# Aviadenovirus structure: a highly thermostable capsid in the absence of stabilizing proteins

**DOI:** 10.1101/2025.04.23.650237

**Authors:** Marta Pérez-Illana, Anna Schachner, Mercedes Hernando-Pérez, Gabriela N. Condezo, Alberto Paradela, Marta Martínez, Roberto Marabini, Michael Hess, Carmen San Martín

## Abstract

High-resolution structural studies have mainly focused on two out of the six adenovirus genera: mastadenoviruses and atadenoviruses. Here we report the high-resolution structure of an aviadenovirus, the poultry pathogen fowl adenovirus serotype 4 (FAdV-C4). FAdV-C4 virions are highly thermostable, despite lacking minor coat and core proteins shown to stabilize the mast- and atadenovirus particles, and having no genus-specific cementing proteins. Unique structural features of the FAdV-C4 hexon include a large insertion at the trimer equatorial region, and a long N-terminal tail. Protein IIIa conformation is closer to atadenoviruses than to mastadenoviruses, while protein VIII diverges from all previously reported structures. We interpret these differences in light of adenovirus evolution. Finally, we discuss the possible role of core composition in determining capsid stability properties. These results enlarge our view on the structural diversity of adenoviruses, and provide useful information to counteract fowl pathogens or use non-human adenoviruses as vectors.

## Introduction

Adenoviruses are found in all types of vertebrates. The International Committee on Taxonomy of Viruses recognizes six genera in the *Adenoviridae* family: mastadenoviruses infecting mammals, aviadenoviruses infecting birds, atadenoviruses found in birds, reptiles and ruminants, siadenoviruses from birds and amphibians, ichtadenoviruses found in fish, and testadenoviruses in turtles (Benkö *et al*., 2022). Adenoviruses stand out among the icosahedral, non-enveloped viruses because of their large capsid size (950 Å, ∼150 MDa), triangulation number (*pseudo*-T = 25) and complex composition. The capsid facets are formed by 240 trimeric hexons, with twelve pentameric pentons at the vertices. Trimeric fibres of variable length and number project away from the vertices [reviewed in (Gallardo *et al*., 2021)]. Minor coat proteins IIIa, VI and VIII on the inner capsid surface contribute to modulate the quasi-equivalent icosahedral interactions, along with the genus specific outer proteins IX (in mastadenoviruses) or LH3 (in atadenoviruses) (Liu *et al*., 2010; Marabini *et al*., 2021; Menéndez-Conejero *et al*., 2017; Natchiar *et al*., 2018; Pantelic *et al*., 2008). The linear dsDNA genome varies in length between genera, ranging from 26 to 49 kbp (Benkö *et al*., 2022; Gallardo *et al*., 2021). The genome is packed within the capsid accompanied with viral “histone-like” proteins (∼25 MDa of protein in the most studied human adenoviruses) (Benevento *et al*., 2014; Hosokawa and Sung, 1976; van Oostrum and Burnett, 1985), in a nucleoproteic structure denominated core. Core proteins V (exclusive from mastadenoviruses), VII and µ regulate genome condensation (Marion *et al*., 2017; Martín-González *et al*., 2019; Martín-González *et al*., 2023; Ortega-Esteban *et al*., 2015b; Vayda *et al*., 1983). The structure of core proteins and their organization within the virion are still not fully resolved (Dai *et al*., 2017; Hernando-Pérez *et al*., 2020; Martín-González *et al*., 2019; Martín-González *et al*., 2023; Pérez-Berná *et al*., 2015; Rafie *et al*., 2021; Schwartz *et al*., 2023).

Adenoviruses are mainly studied for their potential applications in biomedicine, culminating with their use as SARS-CoV-2 vaccine vectors during the COVID-19 pandemic (Hasanpourghadi *et al*., 2021). Simian and low-prevalence human adenoviruses are used as vectors to prevent inactivation by pre-existing immunity, and in this context more exotic types with a phylogenetically more distant origin have also been considered as potential vector candidates. Adenoviruses infecting birds are found in *Aviadenovirus*, *Atadenovirus* and *Siadenovirus* genera. There are currently over 20 recognized aviadenovirus species, of which five infect fowl (FAdV-A to E) (Benkö *et al*., 2022; Schachner *et al*., 2019). Fowl adenoviruses (FAdVs) are pathogens of increasing economic importance in poultry production, with recent decades marked by global emerging of FAdV-associated diseases (Hess, 2000; Schachner *et al*., 2018). Non-pathogenic aviadenoviruses have been proposed as vectors for poultry vaccines and gene delivery, not only for FAdV-caused diseases (Schachner *et al*., 2018), but for other etiological agents as well (Corredor *et al*., 2017). The most remarkable features differentiating aviadenovirus from human adenovirus virions include the longer genome (∼45 *vs* 36 kbp) and the presence of two fibres in each penton, instead of one (Gelderblom and Maichle-Lauppe, 1982; Hess *et al*., 1995). Infectivity assays showed that FAdV-A1 (strain CELO— Chick Embryo Lethal Orphan Virus) particles are highly thermostable compared to the prototype human adenovirus (HAdV-C5) (Michou *et al*., 1999). This observation was intriguing, because aviadenovirus genomes do not code for coat protein IX nor core protein V, both involved in HAdV-C5 capsid stabilization (Bauer *et al*., 2021; Benkö *et al*., 2022; Colby and Shenk, 1981; Martín-González *et al*., 2023; Ugai *et al*., 2007). In atadenoviruses, genus specific proteins LH3, p32k and LH2 have been proposed to substitute proteins IX and V and contribute to increased capsid stabilization (Marabini *et al*., 2021; Menéndez-Conejero *et al*., 2017; Pantelic *et al*., 2008).

High resolution structural studies on adenoviruses have mostly focused on mastadenoviruses: those infecting humans (HAdV-C5, HAdV-D26 and HAdV-F41), ruminants (BAdV-3), or simian adenoviruses used as vectors (ChAdV-Y25) (Baker *et al*., 2021; Cheng *et al*., 2014; Dai *et al*., 2017; Natchiar *et al*., 2018; Pérez-Illana *et al*., 2021b; Rafie *et al*., 2021; Yu *et al*., 2017; Yu *et al*., 2022). So far there is only one available high resolution structure for the complete virion of an adenovirus with non-mammalian host: an atadenovirus infecting lizards, LAdV-2 (Marabini *et al*., 2021). For aviadenoviruses, only crystal structures for isolated virion components, corresponding to the major surface antigens, have been published. These include the hexon and fibre heads of CELO virus (FAdV-A1) (El Bakkouri *et al*., 2008; Guardado-Calvo *et al*., 2007; Xu *et al*., 2007). No structural data, particularly for the entire virion, are available for fowl aviadenoviruses other than the historical prototype. This study provides the first high resolution structure for a complete aviadenovirus virion, FAdV-C4^1^, together with its thermostability properties. The findings shed light on differences with the hitherto known structures of two other adenovirus genera in the context of virus evolution and capsid stabilization.

## Results

### Thermostability of FAdV-C4 virions

We examined the thermostability of FAdV-C4 virions in comparison to HAdV-C5 using extrinsic fluorescence of the DNA-intercalating agent YOYO-1, to characterize capsid disruption as a function of temperature (Hernando-Pérez *et al*., 2020; Pérez-Illana *et al*., 2021a). The fluorescence emission of YOYO-1 increased with temperature for both specimens, as genomes became exposed to the solvent (**Figure 1a**). The half-transition temperature indicating the maximum rate of DNA exposure was T_0.5_ = 57.3 ± 0.01 °C for FAdV-C4, 10 °C higher than for HAdV-C5 (T_0.5_ = 47.5 ± 0.06 °C). In comparison, the highest T_0.5_ reported in human adenoviruses is that of enteric human adenovirus HAdV-F41, with approximately 51°C (Pérez-Illana *et al*., 2021b). Hence, similar to the results previously reported for FAdV-A1 (CELO) (Michou *et al*., 1999), we found that FAdV-C4 virions are highly thermostable compared to human adenoviruses with either respiratory or enteric tropisms.

**Figure 1.**
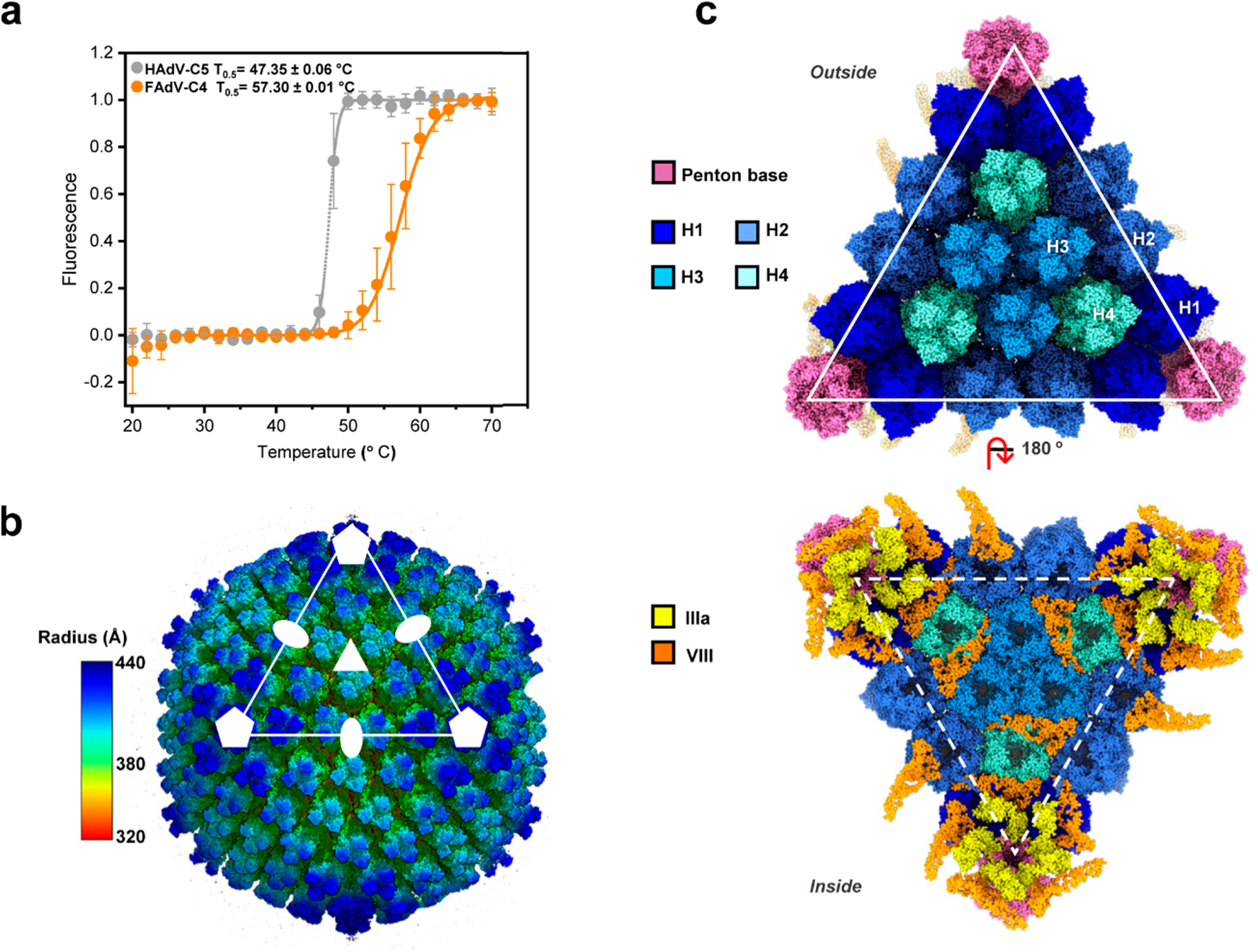
Thermostability and general organization of the FAdV-C4 capsid. **(a)** Effect of heating on HAdV-C5 and FAdV-C4 virions analysed by extrinsic fluorescence. Circles represent the normalized fluorescence ± STD for HAdV-C5 and FAdV-C4. Continuous lines correspond to the Boltzmann sigmoid fitting to estimate the T_0.5_ for each condition tested. **(b)** Radially coloured surface rendering of the 3.3 Å resolution FAdV-C4 map viewed down the 2-fold icosahedral axis. One facet is identified by a white triangle. Icosahedral symmetry axes are indicated with white symbols: 5-fold (pentagon), 3-fold (triangle) and 2-fold (oval). **(c)** Molecular models of the major and minor coat proteins traced in the icosahedral FAdV-C4 facet. Top: view from outside the capsid. Bottom: view from inside the capsid. Protein colours as indicated by the legend at the left-hand side. The four hexon trimers in the AU are numbered H1-H4.

### Absence of genus specific capsid or core proteins in FAdV-C4 virions

We considered the possibility that genus-specific aviadenovirus proteins play a role in capsid stability, as previously reported for mastadenovirus proteins IX and V (Bauer *et al*., 2021; Colby and Shenk, 1981; Martín-González *et al*., 2023; Ugai *et al*., 2007) and atadenovirus protein LH3 (Marabini *et al*., 2021; Menéndez-Conejero *et al*., 2017; Pantelic *et al*., 2008). To investigate this possibility, we analysed the protein composition of purified FAdV-C4 virions by liquid chromatography-tandem mass spectrometry (LC-MS/MS) (**Table S1**). All the expected capsid (hexon, penton base, fibre-1, fibre-2, IIIa, VI, VIII) and core (VII, X) proteins were detected, along with packaging factors IVa2 and L1 52/55 kDa, the terminal protein (TP, with only two copies per virion) and the adenovirus maturation protease (AVP), also packaged together with the genome (Gallardo *et al*., 2021; Mangel and San Martín, 2014). Analysis of tryptic and non-tryptic peptides supports the universality of the AVP cleavage specificity across the *Adenoviridae* family (**Figure S1**). Although FAdVs have extensive left and right ends in their long genomes accommodating unique open reading frames that are genus- and even type-specific (including ORFs 1B, 20A and 28 in FAdV-C4) (Benkö *et al*., 2022; Davison *et al*., 2003; Griffin and Nagy, 2011; Lu *et al*., 2023), we did not detect any peptides corresponding to products of these genes in our assay.

### FAdV-C4 overall structure

We obtained high resolution 3D maps for two different FAdV-C4 strains: KR5, and AG234 (**Table S2, Figure S2a-b**). Model building (**Table S3**) was carried out for the KR5 map (**Figure 1b-c**), given that the AG234 map presented a disrupted particle lacking pentons (**Fig. S2c, Supplementary text**). The structure shows the general characteristics common to all previously solved adenovirus structures: hexons (major capsid protein) and penton base (vertex protein) arranged in a *pseudo* T=25 icosahedral particle, with inner cementing proteins IIIa and VIII on the inner capsid surface (**Figure 1b-c**, **Table S4**). **Table S5** summarizes the main differences between the FAdV-C4 virion proteins and their counterparts in HAdV-C5. Given the known lack of the gene coding for protein IX in aviadenovirus genomes, it was not surprising that we could not trace this protein in the cryo-EM map. In agreement with the LC-MS/MS data, no densities that could be interpreted as genus specific minor coat proteins were present in the map. These results corroborate the lack of genus specific proteins compensating for absence of proteins IX or V in the FAdV-C4 capsid.

### Major coat protein: hexon

As expected, the general architecture of the FAdV-C4 hexon monomer is highly conserved compared to other adenoviruses, with two jelly-rolls (P1 and P2, **Figure 2a**) forming the pseudo-hexagonal base, and flexible loops that contain the hyper variable regions (HVRs) (Rux *et al*., 2003) on the surface-exposed towers (**Figure 2a**).

**Figure 2.**
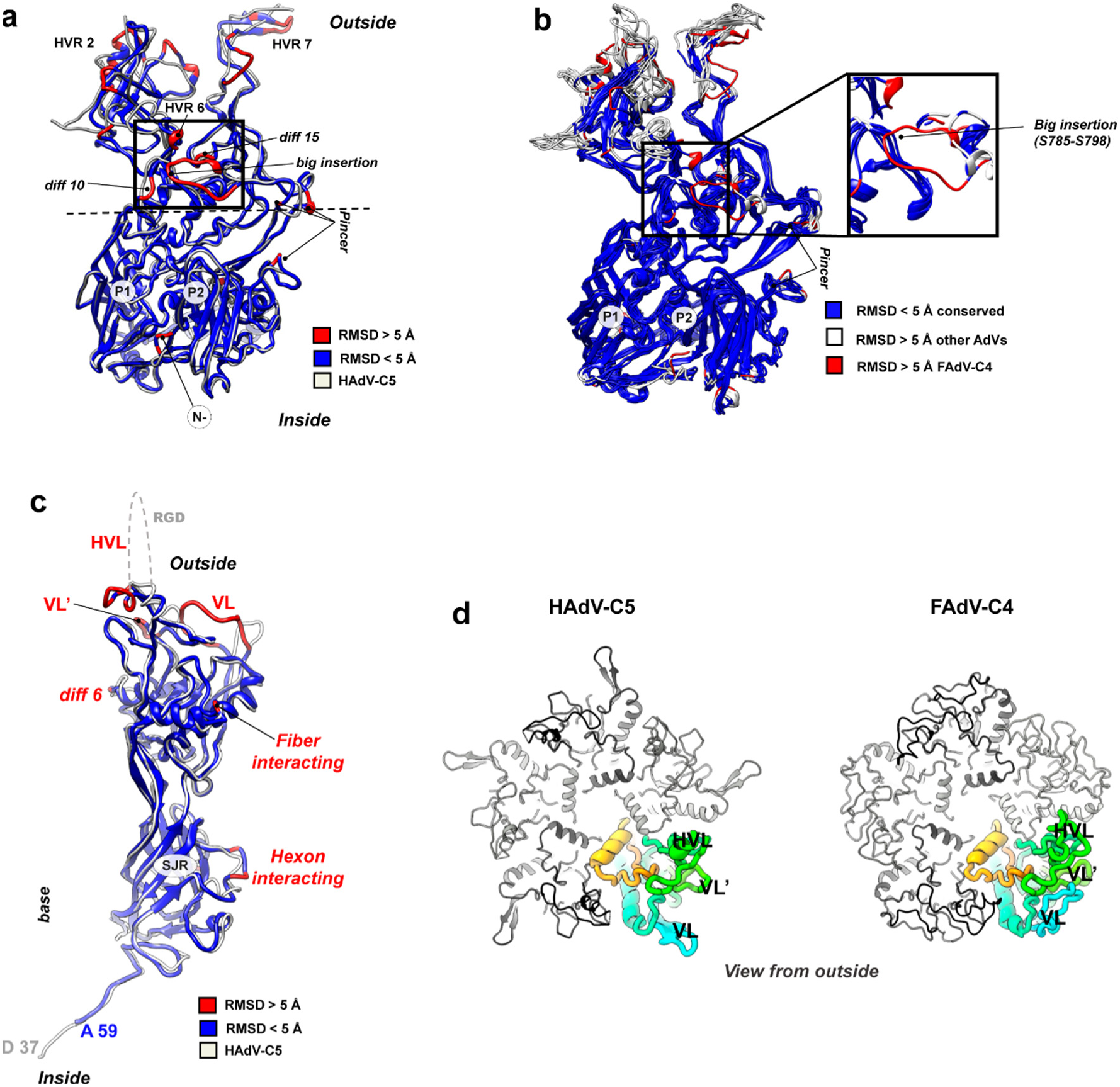
Hexon and penton structures in FAdV-C4. **(a)** Superposition of the HAdV-C5 (PDB ID 6b1t, in white) and FAdV-C4 (coloured by RMSD) hexon monomers. P1 and P2 indicate the two jelly rolls. The horizontal dashed line indicates the equatorial region. **(b)** Structural superposition of available hexon structures (PDB IDs: 6cgv: HAdV-C5, 5tx1: HAdV-D26, 6yba: HAdV-F41, 2obe: SAdV-25, 3zif: BAdV-3, and 6qi5: LAdV-2). Conserved residues (with < 5Å RMSD compared to the rest of structures) are shown in dark blue. Residues with RMSD higher than 5 Å are represented in white for all hexons, except for the >5 Å RMSD FAdV-C4 residues, which are represented in red, to be distinguished from the rest. Conserved residues are mainly forming the jelly rolls in the base. Variable regions are mostly located at the top of the hexons, conforming the specific HVRs. Note the unique big insertion of FAdV-C4 (zoom at the right), not present in any of the other hexon structures. **(c)** Superposition of the HAdV-C5 (PDB ID 6b1t, white) and FAdV-C4 (coloured by RMSD) penton base monomers. The hypervariable and variable loops (HVL, VL and VL’) along with other diff regions are indicated. **(d)** Top view of the penton base pentamers of HAdV-C5 and FAdV-C4, shown with one monomer rainbow-colored and with the HVL, VL and VL’ indicated, and the rest of monomers in grey tones.

Structural superposition of HAdV-C5 and FAdV-C4 hexon monomers revealed seventeen regions of difference (*diff*) with RMSD > 5 Å (**Table S6, Figure 2a**). The N-terminal region of the FAdV-C4 hexon is five amino acids longer (**Table S6**, *diff 1*), a unique feature of aviadenovirus hexons (Xu *et al*., 2007) which could have an impact in the interaction with other proteins (see section “Protein networks in FAdV-C4”). We could fully trace the seven HVRs of FAdV-C4 (**Table S4**), which as expected differ from the corresponding regions in HAdV-C5 (**Table S6**, *diff 2-4* and *6-9*). *Diff 11, 12* and *16* are located in the equatorial region of the hexon monomer, between the base and the tower regions. In the three-dimensional structure, these equatorial differences come together forming a sort of *pincer* above the second jelly roll (**Figure 2a and Table S6, *“pincer*”**). The pincer is located close to the position occupied by the outer cementing protein IX in HAdV-C5 (Liu *et al*., 2010), suggesting that the differences in this region could be related to the absence of outer cementing proteins in FAdV-C4. Indeed, several residues in FAdV-C4 hexon would clash with protein IX if this protein was present in FAdV-C4 (**Figure S3a**). Multiple sequence (Rux *et al*., 2003) and structural alignments indicate that the *pincer* is one of the variable regions when comparing all available hexon structures (**Figure 2b**). HAdV-F41 hexon shows a subtle difference in this area when compared to HAdV-C5, proposed to be involved in the unique conformation of protein IX in the enteric virus (Pérez-Illana *et al*., 2021b). Likewise, LAdV-2 hexon exhibits differences in this region (Marabini *et al*., 2021), and its genus specific cementing protein LH3 also follows a different conformation when compared to that of protein IX in HAdV-C5. These observations highlight the pincer region as an indicator for evolutionary variability in adenoviruses.

Another difference likely related to the absence of protein IX concerns the structure of HVR2. In HAdV-C5, this loop reaches out from the towers towards the neighbouring hexons **(Figure S3b**), and interacts with protein IX C-terminal helix bundle in hexons 2 and 4 across the icosahedron edge (Liu *et al*., 2010). HVR2 in FAdV-C4 does not spread out from the tower, but folds back towards the main body of the hexon trimer **(Figure S3b)**. It has previously been proposed that hexon HVR2 and the *pincer* region jointly modulate the organization of protein IX on the capsid surface (Reddy, 2017).

D*iff 14* corresponds to a *big insertion* (14 residues) in FAdV-C4 hexon (**Table S6**, **Figure 2a**). Multiple comparison of available hexon structures shows that this feature is unique for aviadenoviruses (**Figure 2b**). The *big insertion* forms part of the loops arising from the P2 jelly roll, but it is not located in the hexon towers as the HVRs. Instead, it contours the equatorial region of the monomer surface and reaches towards the loops arising from the P1 jelly roll, and may be the cause of rearrangements in the neighbourhood (*diff 10* and *diff 15*, **Figure 2a**). Additionally, the *big insertion* forms a negatively charged surface patch (**Figure S3b**). Since all contacts established by the *big insertion* are intra-monomer (**Table S7**), its presence is not expected to impact on the stability of the hexon trimer or the interactions between capsomers in the virion. Finally, *diff 13* (**Table S6**) is located on the surface of the inner hexon cavity— Tyr 756 in FAdV-C4, Tyr 787 in HAdV-C5, and might be involved in specific interactions with proteins VI and VII (Dai *et al*., 2017).

We also analysed the differences between the twelve hexon monomers in the FAdV-C4 asymmetric unit (AU). Regions showing a RMSD >2 Å are located at the trimer base and comprise N-terminal and C-terminal residues (**Figure S3c**). In other reported AdV structures the hexon N- and C-termini also exhibit variability, related to the hexon monomer position in the AU and its neighbouring proteins (Liu *et al*., 2010; Marabini *et al*., 2021). Variability among the twelve monomers in the AU at regions such as the HVRs or the jelly roll bases has been observed in other AdV structures (Marabini *et al*., 2021; Yu *et al*., 2017), but not in FAdV-C4 hexons.

### Vertex capsomer: penton base

Penton base topology is conserved when compared with other AdVs, with an extended N-terminal arm preceding a single jelly roll domain at the pentamer base, and a protruding domain containing variable regions exposed on the capsid surface (**Figure 2c**) (Gallardo *et al*., 2021). Nevertheless, structural superposition of HAdV-C5 and FAdV-C4 penton base monomers revealed eight *diff* regions (**Table S8, Figure 2c**). The N-terminal arm of FAdV-C4 is longer (∼65 amino acids) compared to that of mastadenoviruses (approx. 50 aa) and atadenoviruses (approx. 15 aa) (Liu *et al*., 2010; Marabini *et al*., 2021). We were able to start tracing the FAdV-C4 penton base at Ala59, but not the upstream residues traced in HAdV-C5 (Asp37-Gly50) and LAdV-2 (Glu2-Gly16). A conspicuous polyproline stretch (residues 11-16, **Figure S4a**) may impact on the flexibility of the FAdV-C4 N-terminal arm, and therefore its interactions with neighbouring proteins or cell factors during the viral cycle. The variable loop (VL, *diff 3*) is quite long in FAdV-C4 (17 *vs* 6 aa in HAdV-C5) and folds back towards the main body of the monomer, augmenting intramolecular interactions (**Table S9**), in sharp contrast to the protruding orientation of the VL in HAdV-C5 (**Figure 2c-d**). FAdV-C4 lacks the integrin-binding RGD motif, present in the hypervariable loop (HVL) in HAdV-C5 (*diff 4*) (Sheppard and Trist, 1992). The HVL is notably shorter in FAdV-C4 (5 *vs* 74 aa in HAdV-C5) allowing us to completely trace it. *Diff 5* forms another variable loop (VL’) located between the HVL and the VL and three amino acids longer in FAdV-C4 than in HAdV-C5. We hypothesize that the architectural differences in penton base loops VL, HVL and VL’ could be related with the need to attach two fibres to the same penton base (Hess *et al*., 1995). The arrangement of two trimeric fibres bound to the pentameric penton base is one of the most intriguing features of the aviadenovirus capsid, which cannot be elucidated using icosahedrally averaged maps (**Supplementary text and Figure S4**). Finally, *diff 2* and *diff 7*, located at the base domain, contact the peripentonal hexons and may contribute to the specificity of hexon-penton interactions during assembly (Pérez-Illana *et al*., 2021b; Rafie *et al*., 2021).

### Minor coat protein VIII

Based on the mastadenovirus AVP consensus sequence motif (Mangel and San Martín, 2014), protein VIII in FAdV-C4 is predicted to be cleaved at least at four sites during maturation (**Figure S1**). We have traced the two largest fragments (residues 1-115 and 174-241) almost in their entirety (**Table S4 and Figure 3a**). As in all previously solved AdV structures, there are two independent copies of protein VIII per AU: one gluing the GOS (Group of Six: penton base + five surrounding hexons) and the other beneath the GON (Group of Nine: nine hexons in the facet) (**Figure 1c, bottom and Figure S5**) (Liu *et al*., 2010). Both copies are virtually identical.

**Figure 3.**
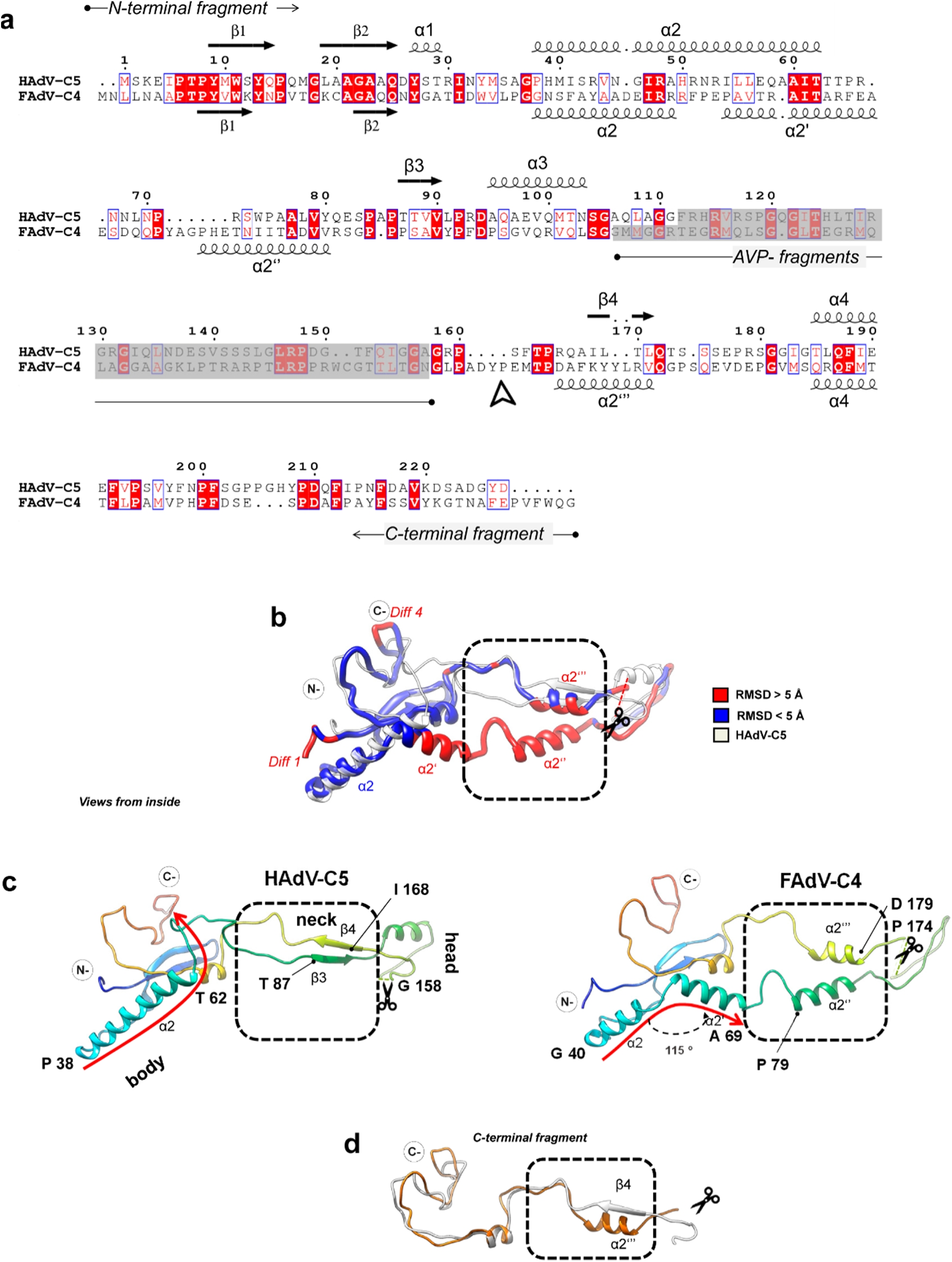
Protein VIII divergent structure in FAdV-C4. **(a)** Sequence alignment and secondary structure elements of protein VIII in HAdV-C5 **(top)** and FAdV-C4 **(bottom)**. Squiggles indicate α-helices and arrows indicate β-strands. Sequence numbering corresponds to HAdV-C5. Residues are coloured according to similarity: black (low similarity), red with blue frame (high similarity) and white on red background (strict identity). N- and C-terminal fragments are delimited by the AVP-processed peptides shadowed in grey. An arrowhead below the FAdV-C4 sequence indicates where the C-terminal fragment tracing starts, seven amino acids downstream of the AVP site. Note the divergence in α2/α2’ and that FAdV-C4 α2’’ and α2’’’ are displaced with respect to β3 and β4 in HAdV-C5. **(b)** Superposition of protein VIII in HAdV-C5 (white, PDB ID: 6b1t) and FAdV-C4 (coloured by RMSD). **(c)** Rainbow coloured structures of protein VIII in HAdV-C5 **(left)** and FAdV-C4 **(right)**. The body, neck (dashed box) and head domains are indicated. Dashed gaps corresponding to the AVP cleaved segment (not traced) are indicated with scissors. Note that α2 in FAdV-C4 is shorter than α2 in HAdV-C5, and along with α2’ follows a different path compared to α2 and subsequent coil structure in HAdV-C5 (curved red arrows). β3 and β4 in HAdV-C5 are distinct from α2’’ and α2’’’ in FAdV-C4 (dashed frame). Helix α2’ in FAdV-C4 is bent by 115 ° with respect to α2. **(d)** Detail of the HAdV-C5 (white) and FAdV-C4 (orange) C-terminal fragment of protein VIII.

Superposition of the HAdV-C5 and FAdV-C4 protein VIII structures reveals several major differences (**Table S10, Figure 3b**). The α2 helix in the body domain of HAdV-C5 protein VIII is 22 amino acid long, while its counterpart in FAdV-C4 is shorter (13 residues) and followed by a bend of 115° and another 12-residue α-helix (α2’) (**Figure 3c**). This bend determines a completely different path for the protein backbone as it leaves the body to enter the neck domain (**Figure 3c, red arrows**). The neck domain, which contains two antiparallel β-strands forming a narrow sheet in HAdV-C5 (β3 and β4), is broader and formed by two helices (α2’’ and α2’’’) in FAdV-C4 (**Figure 3b-c, dashed box**). The head domain is intercalated between these two helices and contains the gap left by the protease cleavage, separating the N-and C-terminal fragments (**Figure 3c, scissors**). The structure of the C-terminal fragment is more conserved and only differs in the secondary structure elements of the neck domain (β4 in HAdV-C5 *vs* α2’’’ in FAdV-C4 (**Figure 3d).** The structure of protein VIII in FAdV-C4 considerably differs from all the previously reported, including those of HAdV-C5, HAdV-D26 and HAdV-F41, and the atadenovirus LAdV-2, where only a small distortion of the neck domain fold was observed (Dai *et al*., 2017; Liu *et al*., 2010; Marabini *et al*., 2021; Pérez-Illana *et al*., 2021b; Rafie *et al*., 2021; Yu *et al*., 2017). This conformational divergence impacts on its interactions with neighbouring proteins (**see section Results. Protein interaction networks in FAdV-C4**).

### Minor coat protein IIIa

There is one copy of IIIa per AU, located beneath the penton **(Figure 1c, bottom).** Similarly to all previously reported adenovirus structures, we were able to trace only 244 amino acids of protein IIIa (out of a total of 590, **Table S4**). From N- to C-termini, the traced structure folds into the same domains previously defined for HAdV-C5: GOS-glue (cementing penton and peripentonal hexons), connecting helix, VIII-binding (interacting with the peripentonal copy of protein VIII), and core proximal (Liu *et al*., 2010). The structural superposition of HAdV-C5 and FAdV-C4 protein IIIa structures reveals several major differences (**Table S11, Figure 4a**). The N-terminus of protein IIIa is ten amino acids shorter in FAdV-C4 than in HAdV-C5 (*diff 1*). Tracing started at Thr16 for FAdV-C4 (**Figure 4a**). In the atadenovirus LAdV-2 this region is also shorter than in HAdV-C5, but its structure is ordered from Gln2 (**Figure 4b**). As a result, in FAdV-C4 the GOS-glue domain shows only one tier of ordered amino acids, compared with the two tiers present in the other solved IIIa structures (**Figure 4a-b, arrowheads)**. Since in other adenoviruses this region interacts with peripentonal hexons (Liu *et al*., 2010; Marabini *et al*., 2021) and the N-terminal arm of penton base (Marabini *et al*., 2021), its lack of order suggests a different network of interactions stabilizing the vertex region in FAdV-C4.

**Figure 4.**
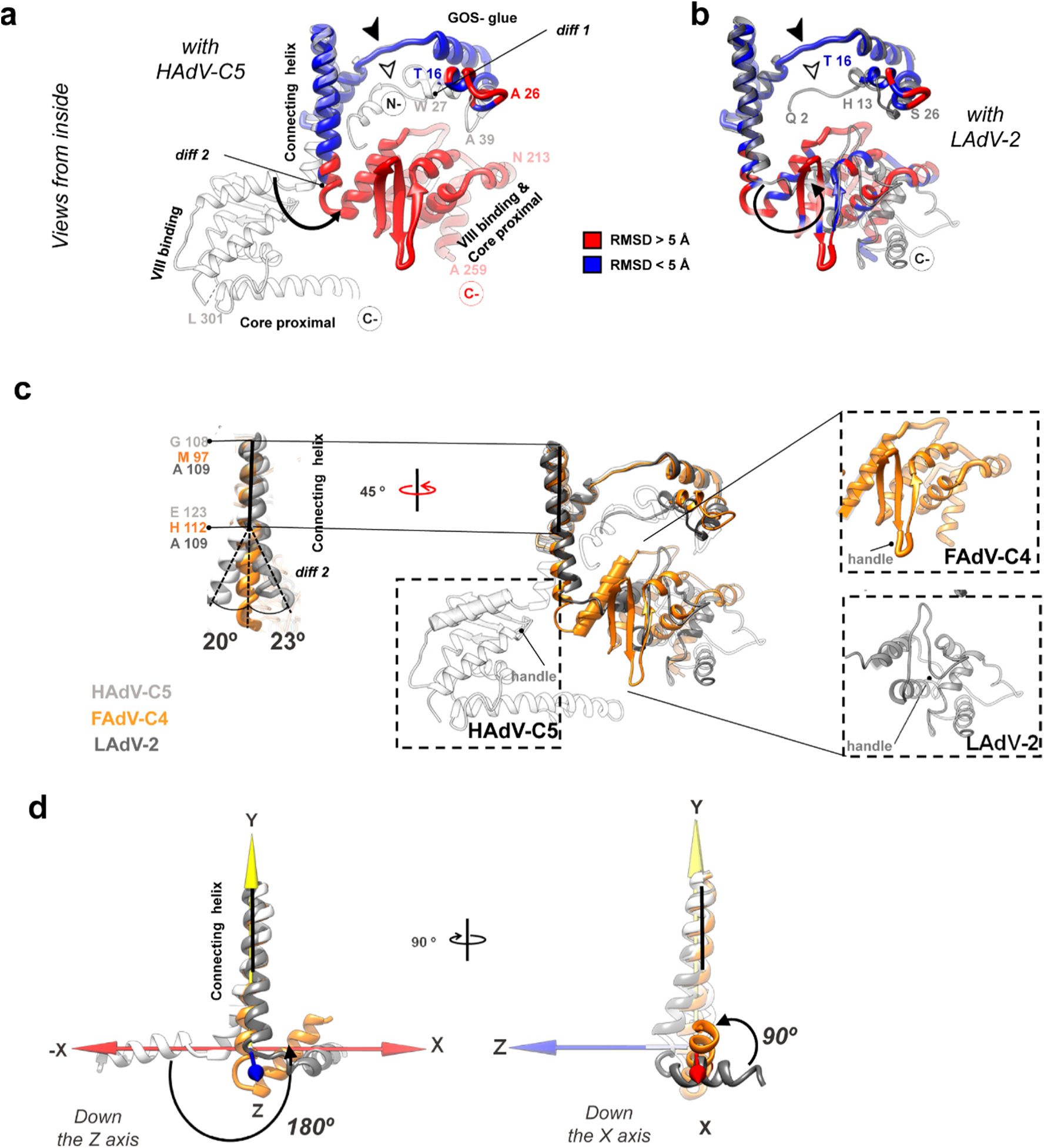
FAdV-C4 minor coat protein IIIa. **(a)** Superposition of protein IIIa in HAdV-C5 (PDB ID: 6b1t, white) and FAdV-C4 (coloured by RMSD), presented in their native position in the capsid. The view is from inside the capsid, with the 5-fold symmetry axis at the top of the page. The GOS-glue, connecting helix, VIII-binding and core proximal domains are indicated. **(b)** As in (a), but comparing with LAdV-2 protein IIIa in grey (PDB ID: 6qi5). The GOS-glue domains and two thirds of the connecting helices mostly overlap in the three structures (FAdV-C4, HAdV-C5 and LAdV-2). The N-terminal region in FAdV-C4 presents fewer ordered amino acids (filled arrowheads), lacking a ∼12 residue stretch ordered in HAdV-C5 and LAdV-2 (hollow arrowheads). The VIII-binding domain is significantly rotated with respect to its position in HAdV-C5 and slightly rotated with respect to LAdV-2. **(c) Centre:** superposition of the three protein IIIa structures: FAdV-C4 (orange), HAdV-C5 (white) and LAdV-2 (grey), represented in their native position in the capsid. **Left:** comparison of the three connecting helices. Horizontal black lines indicate the region where the three helices overlap; dashed lines indicate their bending away from each other (*diff 2*). The dashed black rectangles show the VIII-binding domains in the three viruses, with the small handle formed by *diff3* labelled as a landmark. A cylinder indicates the first helix of the VIII-binding domain in FAdV-C4. **(d)** Focus on the connecting helix and the first helix in the VIII-binding domain showing how their relative orientations vary between the three viruses. For convenience a cartesian coordinate system is represented with the connecting helix of FAdV-C4 along the Y axis and the first helix of the VIII-binding domain of HAdV-C5 along the X axis. The origin of coordinates is located at the last amino acid of the FAdV-C4 connecting helix.

The next point of structural divergence between protein IIIa in the different AdV genera is the C-terminal part of the connecting helix (*diff 2*). The helix in FAdV-C4 is straight, compared to the curved helices in HAdV-C5 and LAdV-2. While the first two thirds of the helix overlap perfectly in the three viruses, at the end the helix in the human and reptilian viruses deviate by *ca.* 20°, each one in a different direction, from the avian one (**Figure 4c, left**). The helix is followed by a flexible loop that, along with the bend of the connecting helix, determines the relative orientation of the VIII-binding domain with respect to the rest of the protein. As a result, the VIII-binding domain is differently oriented in the human, reptilian and avian adenoviruses. In both LAdV-2 and FAdV-C4, the VIII-binding domain swings away from its position in HAdV-C5 by more than 180° (**Figure 4c, center**). Additionally, while in the human and lizard viruses the first helix of the VIII-binding domain is roughly perpendicular to the connecting helix, in FAdV-C4 this helix adopts an oblique orientation (**Figure 4d)**, which results in a completely different conformation of the protein in each one of the genera (**File S1**). This difference in conformation directly impinges on the network of contacts established by protein IIIa in the virion (**see section Results. Protein interaction networks in FAdV-C4**).

In spite of this drastic change in the relative position of domains, the backbone of protein IIIa VIII-binding domain is conserved when compared to the other adenoviruses, except for two small regions (*diff3* and *diff4)* (**Table S11, Fig S6a)** that show variability among the three structures. Finally, the core proximal domain, formed by the last traced residues in HAdV-C5, does not seem to be ordered in FAdV-C4 (*diff 5)* (**Figure S6a**), as similarly reported for the atadenovirus LAdV-2 (Marabini *et al*., 2021).

### Additional internal densities

Analysis of densities that did not allow unequivocal polypeptide tracing (remnant densities, RD) revealed several features of interest (**Figure 5**). By comparison to other AdVs, *RD1*, with three copies located at the mouth of each hexon, presumably corresponds to peptides cleaved during maturation from the precursor forms of proteins VI and VII (Dai *et al*., 2017). L-shaped densities at the local 3-fold symmetry axis between hexons 2, 3 and 4 (*RD2a*), and an equivalent 3-fold averaged density at the icosahedral 3-fold (*RD2b*) have also been observed in human adenovirus maps (Dai *et al*., 2017; Pérez-Illana *et al*., 2021b; Rafie *et al*., 2021; Yu *et al*., 2017). In HAdV-F41, *RD2a* was interpreted as part of core protein V (Rafie *et al*., 2021). However, FAdV-C4 lacks protein V and does not have genus-specific proteins substituting for V. This observation suggests that a virion component distinct from protein V, present in both mastadenoviruses and aviadenoviruses, must be filling this position. The fact that HAdV-C5 particles devoid of protein V also present these *RD2a* densities further support this hypothesis (Martín-González *et al*., 2023).

**Figure 5.**
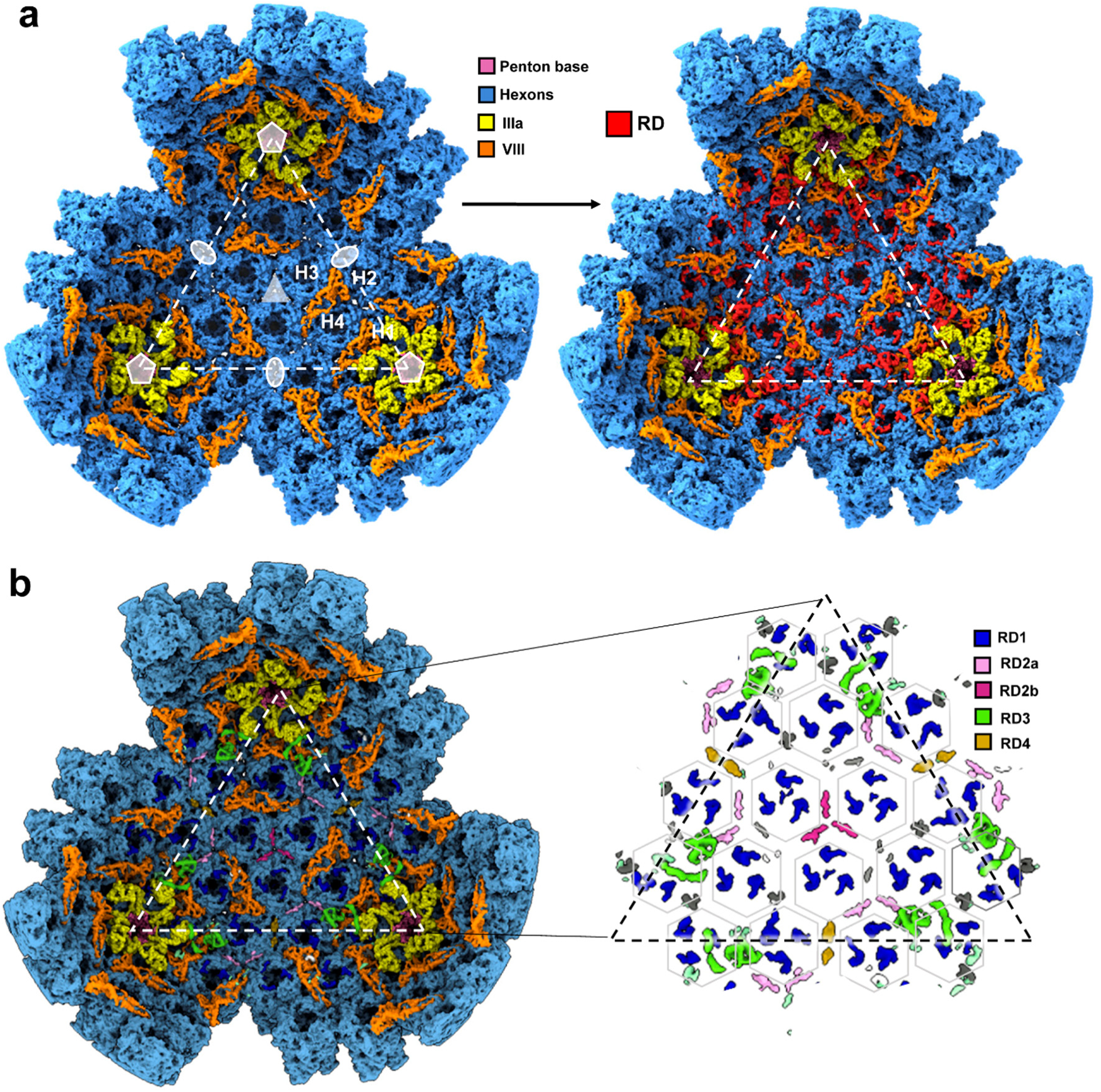
Additional internal components of the FAdV-C4 capsid. **(a)** FAdV-C4 map, as viewed from inside of the capsid. **Left:** only the density corresponding to the traced proteins is shown. The two-fold (oval), three-fold (triangle) and five-fold (pentagon) symmetry axes, the four hexons in one AU (H1-H4), and the facet (dashed triangle) are indicated for reference. **Right:** remnant densities (RD) are coloured in red. **(b)** Remnant map components colour-coded as RD1-4. **Right:** a zoom showing only the colour-coded RDs. RDs considered too small to be analysed are left in grey. Densities correspond to the unsharpened map contoured at 1.5 σ. Hexagon contours indicate hexon positions.

Four α-helices can be modelled in *RD3*, located at the vertex near proteins IIIa and VIII. (**Figure S6b**). This is reminiscent of the appendage domain (APD) of protein IIIa, a region downstream of the core proximal domain observed in HAdV-D26 and also in immature particles of HAdV-C5 (Reddy *et al*., 2022; Yu *et al*., 2017; Yu *et al*., 2022). We tentatively traced amino acids 305-384 of FAdV-C4 protein IIIa in *RD3* (**Table S4**). The topology of the α-helices and the position of *RD3* in the FAdV-C4 capsid do not overlap with those of HAdV-D26, possibly due to the presence of flexible linkers between the helices and to the differently oriented VIII-binding domains. In any case, this *RD3* element would be reinforcing the link between the GOS and the GON. Finally, *RD4* densities are located at the two-fold axes between H2 trimers. Due to the intrinsic curvature of the capsid shell, hexon interactions at the two-fold axes are reduced, compared to the rest of inter-hexon interactions in the capsid (**Table S13**). *RD4* might be contributing to reinforce these weaker hexon-hexon interactions.

### Protein networks in FAdV-C4

Interactions between major and minor capsid proteins have been studied for different adenoviruses, looking for the basis of capsid assembly and stability. Given that FAdV-C4 proteins IIIa and VIII present the largest deviations from the rest of traced capsid components, we focus here on the interactions between these proteins and the rest of the capsid, and how they change with respect to the rest of available AdV structures.

Contacts between neighbouring IIIa monomers are similar in number and location for HAdV-C5 and FAdV-C4, and mostly involve the GOS-glue domain of one monomer and the connecting helix of the other (**Figure 6a, Table S16**). However, the contacts between IIIa and the copy of protein VIII located beneath the GOS diverge noticeably, mostly as a consequence of the large conformational difference found in IIIa. Although the interaction is still established *via* the VIII-binding domain of protein IIIa, the number of contacts decreases by ca. 75% (**Figure 6b, Table S16**). Another consequence of the different conformation of the VIII-binding domain is the loss of interactions between IIIa and hexon 4 (**Figure 6c, Table S16**). Therefore, the change in the orientation of the VIII-binding domain in protein IIIa could in principle weaken the attachment between the GOS and the rest of the capsid.

**Figure 6.**
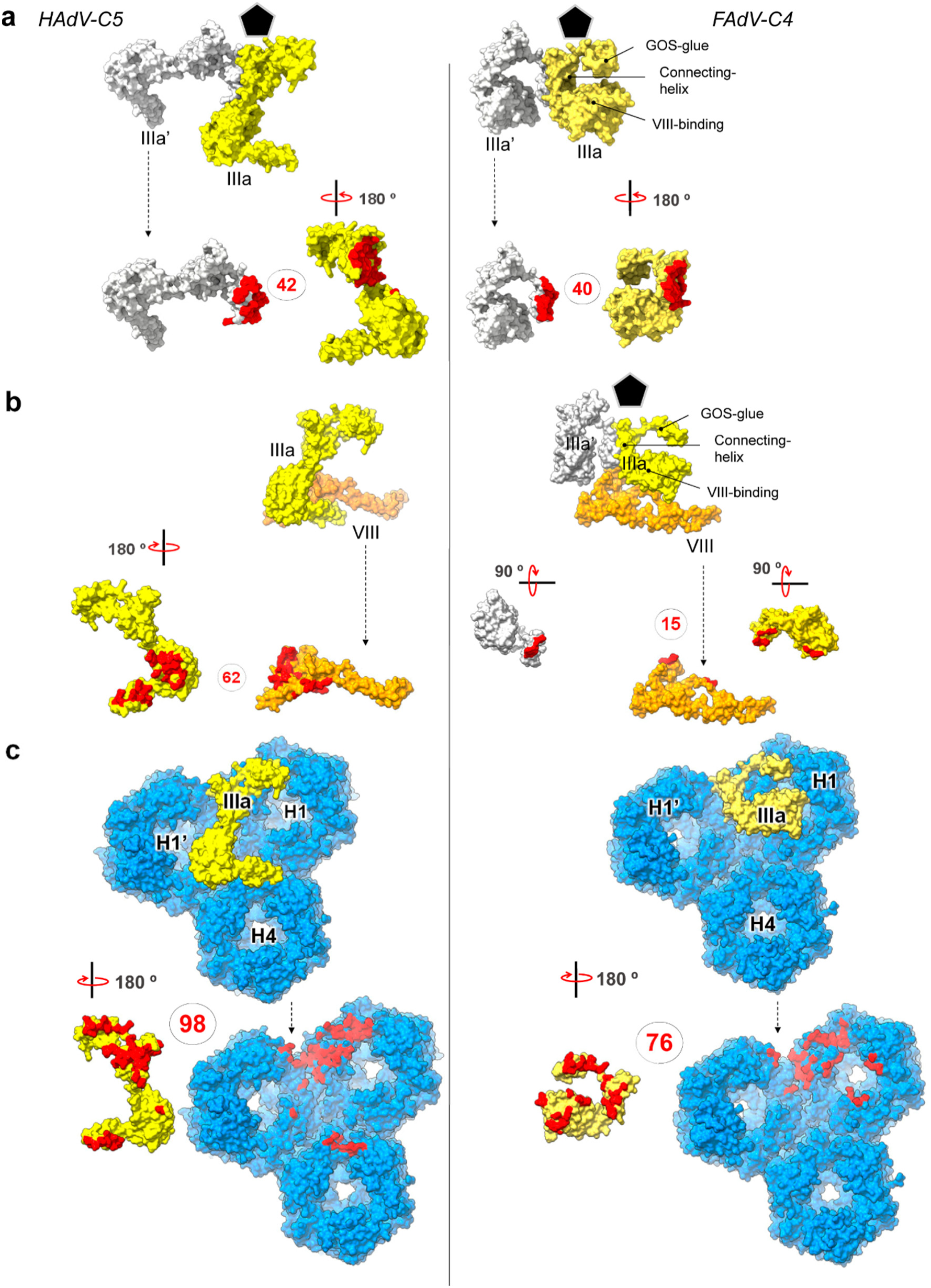
Differences in interactions established by protein IIIa in HAdV-C5 and FAdV-C4. **(a)** Protein-protein interfaces between IIIa (yellow surface) and its neighbouring copy IIIa’ (symmetry copy coloured in white) for HAdV-C5 (left) and FAdV-C4 (right). **(b)** Interfaces between IIIa (yellow surface) and VIII (orange surface). **(c)** Protein-protein interfaces between IIIa (yellow surface) and neighbouring hexons (blue surface). In each panel, the interacting molecules are shown at the top as seen from inside the vertex region (indicated with a black pentagon), and one of them is separated and rotated as indicated at the bottom, to reveal the interfacing residues (coloured in red). Dashed arrows indicate that the molecule orientation is maintained between the top and bottom panels. The number of contacting residues is indicated in red inside circles.

On the other hand, contacts between both copies of protein VIII and the surrounding hexons are slightly more numerous for FAdV-C4 than for HAdV-C5 (**Figure 7a, Table S17**). The “footprint” of VIII on the hexons is broader, reflecting the protein shape. Interestingly, the N-terminal region of one monomer in hexon 1 (chain B) interlaces with the divergent neck region of VIII, while the N-terminus of one monomer in hexon 2 (chain F) crosses over the head domain of protein VIII, further securing it in a clasp near the AVP cleavage site (**Figure 7a-b, ovals, and Figure 6c)**. Thus, the long N-terminal tail unique to the aviadenovirus hexon in combination with the different fold of protein VIII could play a stabilizing role in the capsid. If the structure of protein VIII in FAdV-C4 was the same as in mast- or atadenoviruses, it would clash with the long hexon N-terminal tails.

**Figure 7.**
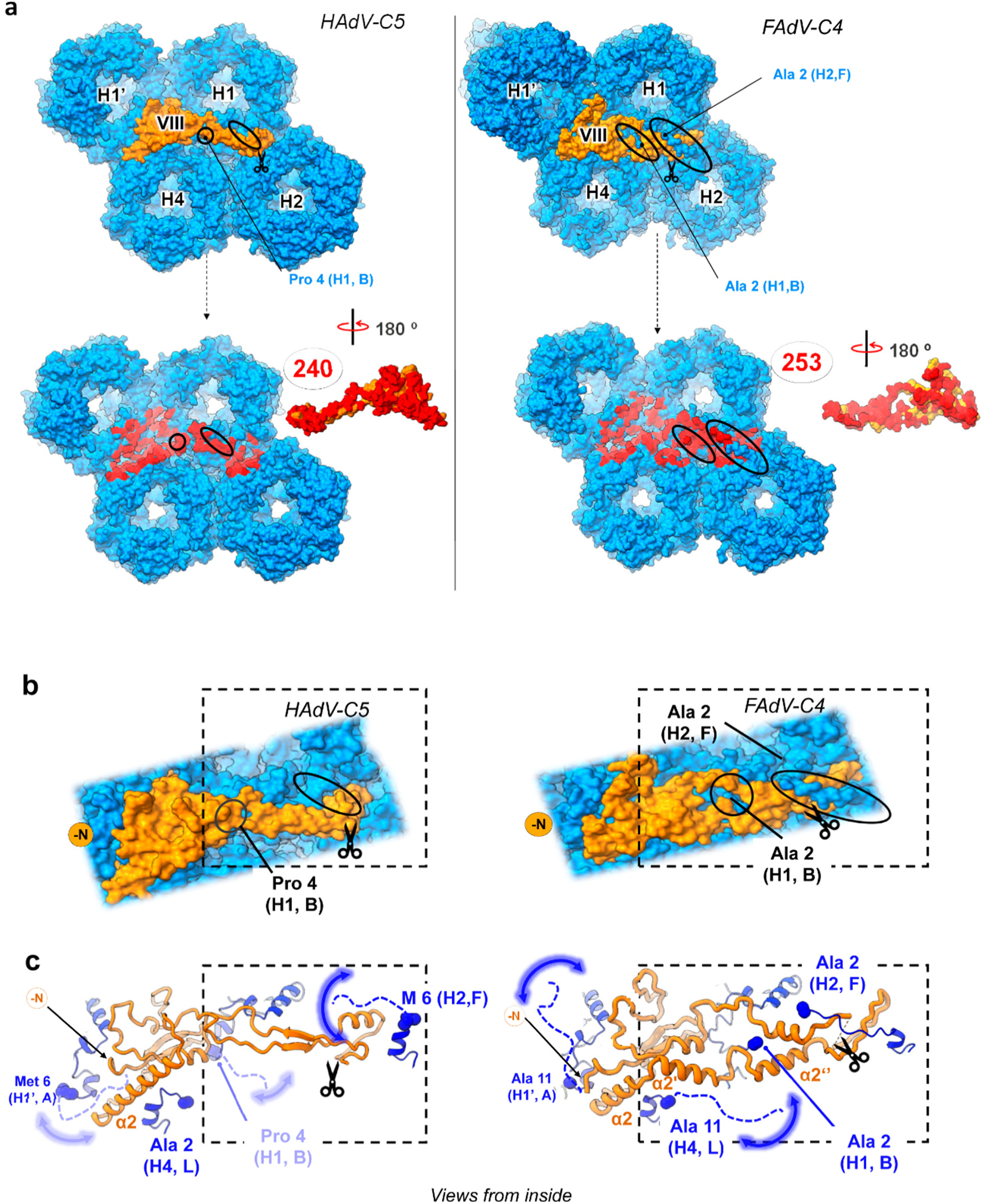
Interfaces between protein VIII and hexons in HAdV-C5 and FAdV-C4. **(a)** Protein-protein interfaces between VIII (orange surface) and hexons (blue surface), with contacting residues highlighted in red, and their number indicated as in Fig. 6. The N-termini of HAdV-C5 hexons are shorter than their FAdV-C4 counterparts (ovals). **(b)** Zoom in on protein VIII. The traced N-termini of the HAdV-C5 hexons (blue) are not interlaced with protein VIII (orange) (left, ovals) whereas the N-termini of the FAdV-C4 hexons (most notably H2) are clasping protein VIII (right, ovals). **(c)** Ribbon representation of the molecules represented as surfaces in (a) and (b), with the non-traced first residues of the hexons indicated by dashed lines. Blurry curved arrows represent the flexibility of the hexon N-termini. Scissors indicate the AVP sites. Dashed frames are drawn to facilitate comparisons. Hexon chain IDs (according to **Table S4**) are indicated in parenthesis.

### Insights into the FAdV-C4 core

Aviadenovirus genomes (including that of FAdV-C4) are among the longest in the *Adenoviridae* family, approximately 25% longer than those of human AdV (approx. 45 kbp *vs* 36 kbp dsDNA) (Benkö *et al*., 2022). So far, a longer genome has only been reported in the single known ichtadenovirus (48 kbp) (Doszpoly *et al*., 2019). All AdV genomes code for core proteins VII and µ, but only mastadenovirus genomes contain the gene for protein V. Although they appear in all AdV genera, proteins VII and µ differ substantially in sequence and length (**Figure 8a**). For example, in mastadenoviruses protein VII is considerably longer than µ (198 *vs* 80 amino acids for the immature polypeptides) while the length relation is reversed in aviadenoviruses (72 residues in VII *vs* 176 aa in µ). In spite of this variability, the N-terminus of pVII and the C-terminal region of protein pre-µ (pµ) are well conserved across all the genera. Remarkably, pVII and pµ in ichtadenoviruses (Doszpoly *et al*., 2019) are reduced to just these two conserved domains. The conserved domains are delimited by consensus maturation cleavage sites (**Figure 8a, dashed lines**). Since AVP is a genus-common gene, proteolytic maturation seems to be an ancient feature of the adenovirus replicative cycle. These observations point towards a hot spot of evolution in the region coding for the core proteins.

**Figure 8.**
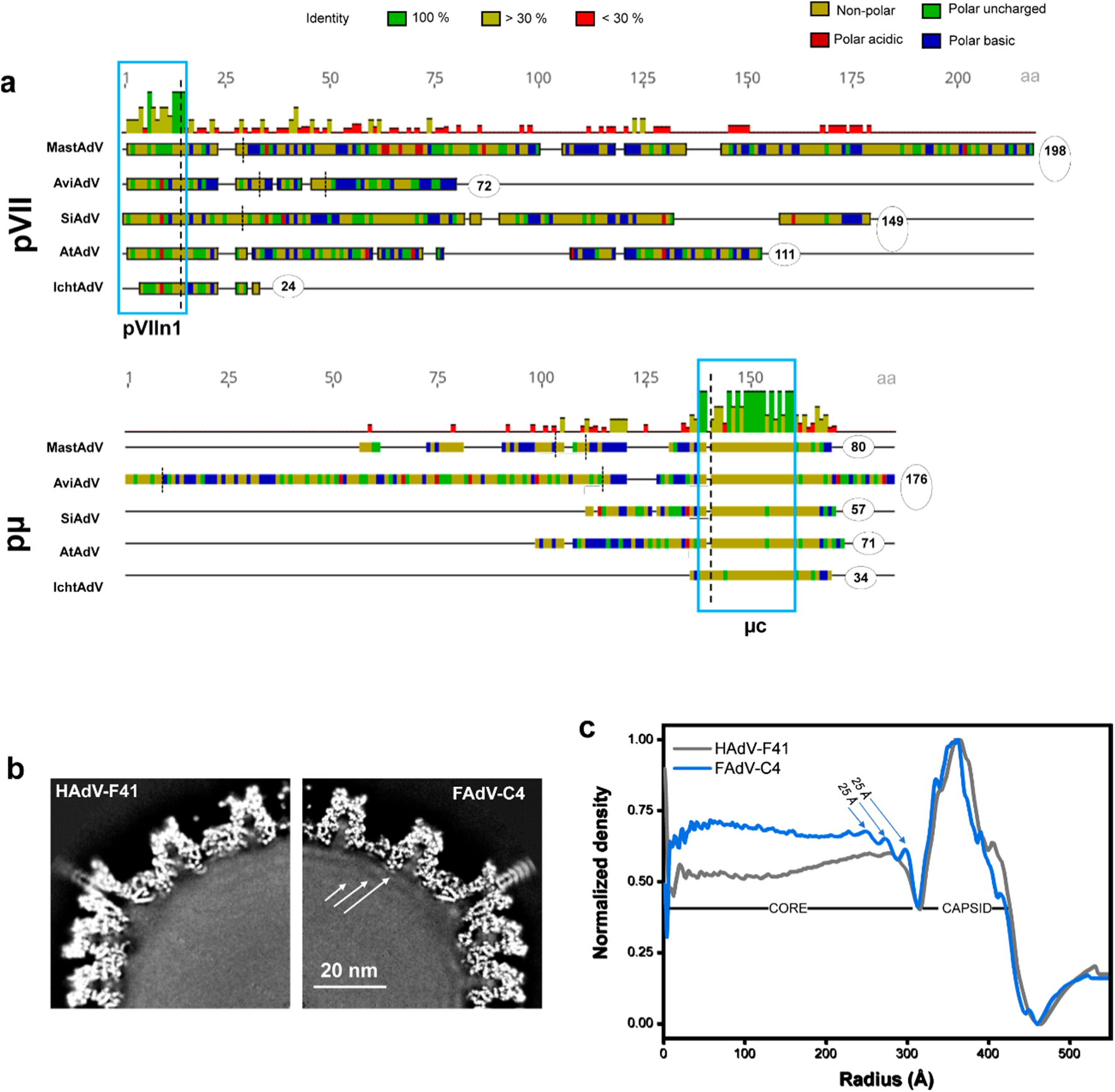
Organization of the core varies among adenovirus genera. **(a)** Multiple protein sequence alignment of core proteins pVII **(top)** and pµ **(bottom)**, showing their respective length (in amino acids) encircled. NCBI GenBank accession numbers for the genomes chosen as representative members of each genus are as follows: MastAdV, HAdV-C5 (AC_000008); AviAdV, FAdV-A1 (AC_000014); SiAdV, FrogAdV-1 (NC_002501); AtAdV, OAdV-7 (NC_004037); IchtAdV, WSAdV (MK_101347). Predicted AVP cleavage sites are indicated by dashed vertical lines following the consensus patterns for proteins pVII and pµ. Amino acids are coloured by polarity, and the top histograms indicate the mean pairwise identity over all pairs in the column. Notice that the number of cleavage sites also varies between genera, but the cuts delimiting the conserved domains of protein VII and µ are also conserved. Blue rectangles highlight high similarity regions corresponding to the pVII N-terminal peptide (pVII_n1_) and C-terminal region of pµ. **(b)** Central section of a HAdV (as an example we use HAdV-F41, EMDB code:10768, **left)** and FAdV-C4 **(right)** cryo-EM maps. Arrows point to more pronounced concentric shells in FAdV-C4. **(c)** Radial average profiles of FAdV-C4 and HAdV-F41 maps. The arrows indicate the 25 Å spacing between shells.

In previously published mastadenovirus (Hernando-Pérez *et al*., 2020; Martín-González *et al*., 2023; Pérez-Berná *et al*., 2009; Pérez-Illana *et al*., 2021b) and atadenovirus (Marabini *et al*., 2021; Menéndez-Conejero *et al*., 2017) cryo-EM maps where the core has not been masked away, there is mostly featureless density inside the capsid. This indicates that the core does not have an icosahedral organization. In FAdV-C4, the unsharpened map presents a few, weak concentric shells near the inner wall of the capsid (**Figure 8b**). The distance between shells is 25 Å (**Figure 8c, arrows**), similar to the distances observed between the concentric layers of naked dsDNA in tailed bacteriophages (Lander *et al*., 2013). The same shells are observed in the FAdV-C4 map lacking pentons (**Figure S2c, d**). The presence of concentric shells suggests that the longer aviadenovirus genome may have a tendency to pack in a more ordered fashion than the mastadenovirus genome. Similar weak shells have been observed in HAdV-C5 particles lacking protein VII, suggesting that they may originate from changes in the DNA/protein balance in the core (Hernando-Pérez *et al*., 2020). Another hint of a different core organization in FAdV-C4 is the occasional observation of material released from disrupted particles in cryo-EM grids, which appears as condensed yarn ball-like objects (**Figure S7**). These are reminiscent of the cores released from immature HAdV-C2 *ts1* particles (Pérez-Berná *et al*., 2009; Pérez-Berná *et al*., 2012) in contrast to cores released by HAdV-C5 virions which show a diffuse aspect. This observation suggests that the mature aviadenovirus core proteins have a stronger DNA condensing action than those in mature mastadenoviruses. Since in HAdV-C2 *ts1* particles the higher core condensation correlates with the higher stability of the immature particle (Ortega-Esteban *et al*., 2013; Ortega-Esteban *et al*., 2015a; Ortega-Esteban *et al*., 2015b; Pérez-Berná *et al*., 2012), this observation prompts the hypothesis that core organization also has a role in the increased thermostability of the aviadenovirus virion. In fact, extrinsic fluorescence analyses show that genome exposure to intercalating dyes requires higher temperatures in FAdV-C4 than in HAdV-C5 not only at neutral pH (**Figure 1a**), but also in a range of acidic conditions (**Figure S8**). Lack of genome accessibility to dyes is also a sign of stronger core condensation (Pérez-Illana *et al*., 2021a).

## Discussion

With the contribution of the first complete fowl aviadenovirus virion structure in this work, there are presently high resolution structures available for three out of the six recognized adenovirus genera: *Mastadenovirus*, *Atadenovirus* (reviewed in (Gallardo *et al*., 2021)), and *Aviadenovirus* (this work). *Adenoviridae* family phylogeny indicates that out of these three genera, aviadenoviruses represent the evolutionarily most distant relatives to the *Mastadenovirus* genus which is the most extensively studied genus to date (Benkö *et al*., 2022). All AdVs share a common general organization of the virion. In particular, hexon and penton base, the major capsid proteins shaping the characteristic icosahedral surface of the virion, show a conserved overall architecture, with the main differences located at surface-exposed loops as a consequence of their interaction with host factors (receptors, immune system). In addition to these expected results, we found that the FAdV-C4 hexon presents a unique insertion of fourteen amino acids located at the equator of the hexon trimer. Contrary to what might be presumed, this insertion was not found to participate in inter-monomer interactions strengthening the trimer. Since it is located on the hexon surface, exposed to the outer part of the capsid, it could be speculated to play a role in virus-host interactions, similar to the hypervariable regions (HVRs) on the towers of HAdV hexons (Khare *et al*., 2012). Thus, the *big insertion* could account for antigenic and/or receptor binding regions specific to fowl adenoviruses. In any case, observation of this site argues for its further investigation, for instance as a candidate for epitope display for the design of fowl adenovirus-based vectors (Hansra *et al*., 2015). In addition, engineering the long equatorial hexon insertion may be facilitated by the lack of external cementing proteins in aviadenoviruses, reducing the likelihood of structural interferences at this site. However, the location of the insertion is not as surface-exposed as the HVRs, which may preclude its interaction with cellular factors. A further candidate site for antigen display was identified in the penton base variable loop VĹ, which is longer in FAdV-C4 compared to HAdV-C5. Similarly, the HVL and VL have already been used for introducing changes into the penton base of HAdVs (Besson *et al*., 2020). Penton base loops, however, are less solvent-exposed in FAdV-C4 than in HAdV, as they fold back onto the main body of the pentamer. This structural peculiarity is noteworthy as it could serve as a mechanism that helps secure the characteristic double fibre of aviadenoviruses.

The results presented here, together with previously reported structures, define three points of variability in the inner cementing protein, IIIa, that enable the diverse conformations found across the members of the *Adenoviridae* family. First, the N-termini show different ordered lengths. Second, the helix connecting the GOS-glue and the VIII-binding domains can bend in different directions, resulting in divergent orientations of the VIII-binding domain. Last, the core proximal and APD domains are also different: while the backbone of each of the domains is fairly conserved among the reported structures, the boundaries and connecting regions are diverse, resulting in a different orientation of the domains in the context of the virion. Hence, protein IIIa is organized in modules connected by flexible hinges. When the structure of LAdV-2 was solved, it was proposed that the difference in IIIa conformation was due to the presence of clashing, genus specific proteins beneath the vertex (Marabini *et al*., 2021). The fact that we did not detect any such proteins in FAdV-C4, where the conformation of IIIa is closer to that of LAdV-2 than to the HAdVs, argues against that hypothesis. Instead, we hypothesize that the architecture of IIIa in avi- and atadenoviruses may be the ancestral one, and the protein had to adopt a new geometry in the course of evolution. This presumed conformational adaptation may be linked to the incorporation of core protein V to the mastadenovirus genomes, possibly requiring some reorganization of the internal virion components.

Our study also adds new findings for protein VIII, a virion component that has so far not been recognized as a source of variability in the organization of AdV capsids. The conformation of protein VIII in FAdV-C4 is markedly different from those observed in other adenovirus genera. An intuitive hypothesis could be that changes in protein VIII are explained by the distinct conformation of protein IIIa in FAdV-C4, this could result in a change of protein VIII. However, protein VIII in LAdV-2 is quite similar to protein VIII of HAdV-C5 (Marabini *et al*., 2021), although LAdV-2 IIIa is more similar to that of FAdV-C4. Moreover, the two copies of protein VIII in the AU show an identical fold, although only one of them is in contact with protein IIIa. An alternative possibility is that the hexon N-termini, markedly longer in FAdV-C4 than in the other adenoviruses, might be guiding the fold of protein VIII. Altogether, our results indicate that adenovirus internal minor coat proteins have genus specific conformations that may be involved in specific interactions for assembly, or provide specific stability properties to the capsid in each virus. Intriguingly, proteins IIIa and VIII also present significant rearrangements when comparing the mature and immature forms of HAdV-C5 (Yu *et al*., 2022). Since conformational plasticity may be required in both processes, virus evolution and virus assembly alike, we conclude that this is an important property of AdV cementing proteins.

Here we show that FAdV-C4 particles are much more thermostable than those of HAdV-C5, withstanding temperatures up to 10°C higher before exposing their genomes to the solvent. This enhanced stability could be related to the host high body temperature (42°C in chickens) or to the need to resist harsh chemical conditions since, along with vertical transmission, FAdVs spread by faecal-oral transmission. The exact molecular basis of virus capsid stability is difficult to unravel, and is still a subject of intense investigation even for the so-called simple viruses (Mateu, 2013). The FAdV-C4 structure described here shows that, as a result of the conformational variability of proteins VIII and IIIa, the interfaces between minor coat proteins, and between minor and major coat proteins on the inner capsid surface, are considerably different from those in HAdV-C5. But it is difficult to conclude if these differences will have an effect in capsid stability, due to the large complexity and sheer number of interactions present in AdV virions. Counterintuitively, the different conformation of protein IIIa in FAdV-C4 causes a reduction in the number of contacts with the neighbouring proteins, which in principle would make FAdV-C4 virions less stable. Likewise, the remarkable absence of external cementing proteins similar to IX in mastadenoviruses or LH3 in atadenoviruses, whose extensive contacts with the surrounding hexons contribute to stabilize the capsids (Colby and Shenk, 1981; Marabini *et al*., 2021; Menéndez-Conejero *et al*., 2017; Pantelic *et al*., 2008), would point to lower stability in aviadenoviruses. On the other hand, protein VIII in FAdV-C4 has more contacts with hexons and interlaces with the long hexon N-termini (**Figure 7**), suggesting a role for VIII-hexon interactions in stabilization. Remnant densities on the inner surface of the icosahedral shell, whose identity is hard to establish, may also play a role in increasing the aviadenovirus resistance to physicochemical stress.

Not only are aviadenovirus capsids highly thermostable in the absence of external stabilizing proteins, but remarkably, they have a higher DNA packaging capacity than all other genera with structures solved. Despite having a 25% longer genome than HAdV-C5, FAdV-C4 lacks protein V and does not have any other protein that could provide positive charges to compensate the extra negative charges of the longer DNA. On the contrary, atadenoviruses have shorter genomes but code for several genus specific proteins that are positively charged and could be involved in DNA negative charge screening, such as p32k or LH2 (Marabini *et al*., 2021; Menéndez-Conejero *et al*., 2017; Pantelic *et al*., 2008). Capsid stability in HAdV-C5 has been previously related with the core state. Maturation of proteins VII and µ, lack of protein VII, or pH neutralization, result in genome decondensation, increase of internal pressure and decreased stability (Hernando-Pérez *et al*., 2020; Martín-González *et al*., 2019; Ortega-Esteban *et al*., 2013; Ortega-Esteban *et al*., 2015b; Pérez-Berná *et al*., 2012; Pérez-Illana *et al*., 2021a). The need to accommodate a longer DNA molecule inside a shell of the same size suggests that core proteins in FAdV-C4 have a stronger condensing power than those of HAdV-C5, to compensate for the higher internal pressure caused by repulsion among the DNA negative charges. This hypothesis is consistent with the observation of condensed cores released from broken capsids in cryo-EM micrographs. Differences in sequence length and composition, as well as lack of data for core protein copy numbers in aviadenoviruses, hinder any further conclusions on the FAdV-C4 core organization and its possible effect on virion stability. Further experimental studies aimed at estimating virion internal pressure, as well as copy numbers and DNA condensing power of core proteins are needed to elucidate the variability of core organization and its role in the infectious cycle of different adenovirus genera.

## Materials and Methods

### Virus production and purification

Two very similar strains (98.1 % global pairwise nucleotide identity) of FAdV-C4 were used. Strain KR5 (Hess, 2000; Kawamura *et al*., 1964; Marek *et al*., 2012) [NCBI GenBank: HE608152.1] was used for the extrinsic fluorescence experiments, mass spectrometry analysis and cryo-EM structure determination of the complete FAdV-C4 virion. KR5 was propagated in primary CEL (chicken embryo liver) cells infected at a multiplicity of infection (MOI) = 1 and harvested at 67 hours post infection (hpi). Strain AG234 (Schachner *et al*., 2019; Schonewille *et al*., 2008) [NCBI GenBank: MK572849.1] yielded a penton-less cryo-EM map and was propagated in LMH (ATCC: CRL-2117, Leghorn Male hepatocellular carcinoma from chicken epithelial liver) cells infected at a MOI = 4 and harvested at 27 hpi. The HAdV-C5/*attP* variant described in (Alba *et al*., 2007) was propagated as previously reported (Condezo *et al*., 2015; Hernando-Pérez *et al*., 2020) and used as a control for extrinsic fluorescence experiments.

For both FAdV-C4 and HAdV-C5, infected cells were centrifuged in a Heraeus 1.0R megafuge for 40 min at 4000 rpm and 4°C, resuspended in 42 ml of medium (from the supernatant) and lysed by four freeze-thaw cycles. Cell lysates were clarified to remove cellular debris by centrifugation in a Heraeus 1.0R megafuge at 4000 rpm for 30 min at 4°C. For double CsCl gradient ultracentrifugation, the supernatant was distributed in 6 tubes containing a discontinuous gradient of 1.25 g/ml and 1.40 g/ml CsCl in TD1X buffer (137 mM NaCl, 5.1 mM KCl, 0.7 mM Na_2_HPO_4_.7H_2_O, 25 mM Tris-HCl pH 7.4) (2.5 ml of each CsCl buffer and 7 ml of supernatant) and centrifuged at 35700 rpm for 90 min at 18°C in a Beckman Optima L-100 XP ultracentrifuge using a Beckman SW41Ti swinging bucket rotor. The low and high density virus bands from each tube were collected (∼1 ml of band by tube) and independently pooled. In the second gradient, the low and high density pooled materials (6 ml) were laid onto 6 ml of a CsCl 1.31 g/ml solution in TD1X buffer and centrifuged for 18 hours at 35700 rpm and 18°C in a Beckman SW41Ti swinging bucket rotor. Bands containing viral particles from each tube were collected and buffer exchanged into HBS (20 mM HEPES, 0.15M NaCl pH7.8) using dialysis filters (10 kDa cut off, Sigma #D9277) for FAdV-C4, or Econo-Pac 10DG disposable chromatography columns (Biorad Cat#732-2010) with molecular weight cutoff of 6000 Dalton for HAdV-C5. Samples were stored at −80°C after adding glycerol to a final concentration of 10%.

### Quantification of Physical Viral Particles

Capsid protein concentration was quantified using hexon fluorescence emission spectra obtained in a Hitachi Model F-2500 FL spectrophotometer. Sample volumes of 0.150 ml were examined in sealed quartz cuvettes. The sample was excited at 285 nm, and the emission was monitored from 310 to 375 nm using excitation and emission slit widths of 10 nm. The spectra were corrected by subtraction of the buffer spectrum. The maximum emission intensity for each spectrum was found at 333 nm and recorded. The concentration (in viral particles per ml, vp/ml) was determined from a calibration curve calculated from a sample with known concentration.

### Extrinsic fluorescence thermostability assays

Capsid thermostability was analysed by measuring extrinsic fluorescence spectra in the presence of an intercalating dye (YOYO-1), thereby detecting accessibility of the virus genome to the medium. The data for HAdV-C5 have been reported previously (Hernando-Pérez *et al*., 2020; Pérez-Illana *et al*., 2021b). The data for FAdV-C4 were collected in the same time period, as part of a large study comparing the thermostability of different adenovirus specimens. Stock virus samples (∼10^12^ vp/ml) of HAdV-C5 as control or FAdV-C4 were diluted to a final concentration of 5×10^9^ particles/ml in a final volume of 800 µl of Sodium-Potassium Phosphate buffer (8 mM Na_2_HPO_4_, 2 mM KH_2_PO_4_, 150 mM NaCl, and 0.1 mM EDTA, pH 7.4) and incubated overnight at 4 °C at 300 rpm shaking. To analyse thermostability in acidic conditions, samples were incubated in citric acid-sodium citrate buffers: 9.5 mM C_6_H_8_O_7,_ 41.5 mM C_6_H_5_O_7_Na_3_.2H_2_O at pH 6; 20.5 mM C_6_H_8_O_7_, 29.5 mM C_6_H_5_O_7_Na_3_.2H_2_O at pH 5, and 33 mM C_6_H_8_O_7_, 17 Mm C_6_H_5_O_7_Na_3_.2H_2_O at pH 4; all with 150 mM NaCl and 0.1 mM EDTA.

A volume of 8.8 μl of YOYO-1 diluted 1:100 in milliQ water from the initial stock (Thermo Fisher Scientific, stock composition: 1 mM YOYO-1 in DMSO) was added to the virus sample and equilibrated for 5 min at 20°C (or 30°C for the experiments in acidic conditions) before data acquisition. Fluorescence emission spectra were obtained employing a Horiba FluoroLog spectrophotometer equipped with a Peltier temperature control device and continuous shaking. The dye was excited at 490 nm wavelength, and fluorescence emission was monitored from 500 to 580 nm using an excitation slit of 2 nm and an emission slit of 5 nm, integration time 2 s and 950 V. Maximal emission intensity occurred at 509 nm. The temperature in the sample cell was increased from 20°C to 70°C for neutral pH assays; for acidic pH assays, temperatures ranged from 30 to 70°C for HAdV-C5, or to 90°C for FAdV-C4. Emission was recorded every 2°C after holding for 0.5 min for equilibration. Each run took approximately one hour and a half to be completed. The fractional change in fluorescence (normalized fluorescence, *F_n_ (T)*), was calculated according to equation (1).

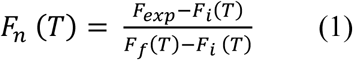

where T indicates the independent variable temperature (°C) and F_exp_ refers to the raw fluorescence signal in counts per second (c.p.s.) at each temperature. F_i_ and F_f_ represent the linear extrapolates, at this temperature, of pre-transition and post-transition base lines, respectively. Buffer signal was almost negligible and was not subtracted.

For assays at neutral pH, a total of six (HAdV-C5) and five (FAdV-C4) independent experiments were considered for each averaged curve, normalized separately according to equation (1) and averaged to obtain the average curve for each condition. Two independent experiments for each virus were carried out for thermostability assays in acidic conditions. Normalized fluorescence *F_n_ (T)* curves were plotted as average values ± standard deviation (STD). The half transition temperatures (*T_0.5_*) were calculated from the fitting of the averaged fluorescence change fraction (*F_n_*) as a function of temperature (*T*) to a Boltzmann sigmoid (Origin software package) according to equation (2):

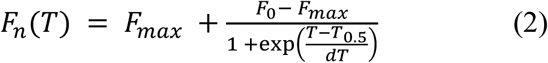

Where *F_0_* ≅0, *F_max_*≅ 1 and *dT* = steepness of the curve. *T_0.5_* are expressed as fit values of each averaged curve ± standard error (SE).

### Mass Spectrometry Analyses

The proteins present in purified FAdV-C4 virions were identified by shotgun proteomic analysis. Briefly, the sample was reduced with 10 mM DTT and alquilated with 55 mM iodoacetamide. To digest the sample, trypsin was added in a 1:25 enzyme:protein ratio (w/w) and incubated overnight. Interfering salts were washed using C18 Zip Tips (Millipore) and the eluting peptides were dried prior to analysis by mass spectrometry. LC-MSMS analysis was performed using an Eksigent 1D-nanoHPLC coupled to a 5600 TripleTOF QTOF mass spectrometer (Sciex, Framinghan, MA, USA). The analytical column used was a C18 Waters nanoACQUITY UPLC 75 µm × 15 cm, 1.7 µm particle size. The trap column was an Acclaim PepMap 100, 5 µm particle diameter, 100 Å pore size, switched on-line with the analytical column. The loading pump delivered a solution of 0.1% formic acid in 98% water / 2% acetonitrile (Scharlab, Barcelona, Spain) at 3 µL/min. The nanopump provided a flow-rate of 250 nL/min and was operated under gradient elution conditions, using 0.1% formic acid (Fluka, Buchs, Switzerland) in water as mobile phase A, and 0.1% formic acid in 100% acetonitrile as mobile phase B. Gradient elution was performed according the following scheme: isocratic conditions of 96% A: 4% B for five minutes, a linear increase to 40% B in 25 min, a linear increase to 95% B in two minutes, isocratic conditions of 95% B for five minutes and return to initial conditions in 10 min. Injection volume was 5 µL. The LC system was coupled *via* a nanospray source to the mass spectrometer. Automatic data-dependent acquisition using dynamic exclusion allowed obtaining both full scan (m/z 350-1250) MS spectra followed by tandem MS CID spectra of the 15 most abundant ions. Acquisition time was 250 ms and 100 ms for MS and MSMS spectra, respectively. MS and MS/MS data were used to search against a customized database containing all the proteins encoded by the FAdV-C4 strain KR5 genome [NCBI GenBank: HE608152.1] and a collection of typical laboratory protein contaminants. Database searches were done using a licensed version of Mascot v.2.6 (Perkins *et al*., 1999), and search parameters were set as follows: carbamidomethyl cysteine as fixed modification and acetyl (protein N-term), NQ deamidation, Glu to pyro-glutamic and oxidized methionine as variable ones. Peptide mass tolerance was set at 25 ppm and 0.1 Da for MS and MS/MS spectra, respectively, and 2 missed cleavages were allowed. Restriction enzyme was set to none. Peptides with individual Mascot ions scores indicating identity or extensive homology (p<0.05) were used for protein identification. Only those proteins with at least one unique peptide were considered. The software Skyline v22.2 (MacLean *et al*., 2010) was used to extract the signal corresponding to the identified protein-specific peptides. The area (arbitrary units) corresponding to each peptide was integrated and summed at the protein level.

### Cryo-EM sample preparation and data collection

*FAdV-C4 strain AG234.* Five hundred µl of 5.6×10^11^ vp/ml FAdV-C4 AG234 purified sample were dialyzed for 1 h at 4°C against phosphate buffered saline (PBS, 137 mM NaCl, 2.7 mM KCl, 10 mM Na_2_HPO_4_, 1.8 mM KH_2_PO_4_ at pH 7.4) and concentrated to 9×10^12^ vp/ml by spinning at 4°C in a 100000 MWCO Amicon Ultra centrifugal filter (Millipore). Particle concentration was further increased by consecutively incubating the grid (Quantifoil R2/4 300 mesh Cu/Rh, glow discharged) on two drops of 1.5 µl of sample (Snijder *et al*., 2017) before the final blotting and plunging in liquid ethane.

*FAdV-C4 strain KR5.* After the first dataset collected (AG234) unexpectedly yielded a map lacking pentons, grids were prepared under different buffer conditions, and a low resolution 3D map was calculated from each one using data collected at the CNB-CSIC cryo-EM 200 kV Talos Arctica, to assess particle concentration and integrity (e.g. presence or absence of pentons). For the data used in the final 3D map, 100 µl of 3.2×10^12^ vp/ml FAdV-C4 KR5 purified sample were dialyzed against 50 mM Tris-HCl pH 7.8, 150 mM NaCl, 10 mM MgCl_2_ for 1 hour at 4°C. Particle concentration was increased by consecutively incubating the grid (Quantifoil R2/2 300 mesh Cu/Rh, glow discharged) on six drops of 3 µl of sample (Snijder *et al*., 2017) before the final blotting and plunging in liquid ethane.

Movies used for high resolution map calculation were recorded at the Diamond Light Source facility (Oxford, United Kingdom) using a 300 kV Titan Krios microscope equipped with a Falcon III detector, with the imaging parameters indicated in Table S2.

### Cryo-EM image processing

All image processing and 3D reconstruction tasks (**Table S2**) were performed within the Scipion framework (de la Rosa-Trevin *et al*., 2016). Frames were aligned using Motioncor2 and weighted according to the electron dose received before averaging (Zheng *et al*., 2017). The contrast transfer function (CTF) was estimated using gCTF (Zhang, 2016). Particles were semi-automatically picked from micrographs corrected for the phase oscillations of the CTF (phase-flipped), extracted into 758 × 758 (FAdV-C4 KR5) or 752 x 752 (FAdV-C4 AG234) pixel boxes, normalized and downsampled by a factor of 2 using Xmipp (de la Rosa-Trevin *et al*., 2013). All 2D and 3D classifications and refinements were performed using RELION (Scheres, 2012). 2D classification was used to discard low quality particles, and run for 25 iterations, with 100 classes, angular sampling 5 and regularization parameter T = 2. Classification in 3D was run for 10 iterations, with three classes, starting with an angular sampling of 3.7° and sequentially decreasing to 0.5, and regularization parameter T = 4. Icosahedral symmetry was imposed throughout the refinement process. The initial reference for 3D classification was a lizard adenovirus cryo-EM map (Marabini *et al*., 2021), low-pass filtered to 60 Å resolution. The class yielding the best resolution was individually refined using the original particles and the map obtained during the 3D classification as a reference, yielding final maps at 3.3 Å and 3.2 Å resolution for FAdV-C4 KR5 and FAdV-C4 AG234 respectively. Resolution was estimated according to the gold-standard FSC = 0.143 criterion as implemented in RELION auto-refine and postprocess routines (Chen *et al*., 2013). The actual sampling for the map was estimated by comparison with a HAdV-C5 model (PDB ID 6b1t) (Dai *et al*., 2017) in UCSF Chimera (Pettersen *et al*., 2004).

Grey values of the maps were standardized with UCSF ChimeraX (Goddard *et al*., 2018) (*volume scale shift-mean factor 1/Standard Deviation*) to have an average = 0 and standard deviation σ = 1 within a region comprising the hexon 1 trimer. Map density thresholds are given as multiples of σ. An Xmipp protocol implemented in Scipion was used to calculate radial average profiles, normalized to minimum and maximum values in the [0, 1] interval.

### Model building and structure analysis

Interpretation of the FAdV-C4 KR5 3D map was performed using the molecular modelling workflow based on sequence homology in Scipion (Martínez *et al*., 2020). The initial model for each polypeptide chain was predicted with Modeller (Webb and Sali, 2016), using as input template the structure of the respective HAdV-C5 homolog chain (PDB ID 6b1t). UCSF Chimera was used to fit each chain initial model as a rigid body into the sharpened map. Next, the fitted model of each chain was iteratively refined using Coot (Emsley *et al*., 2010) and Phenix real space refine (Afonine *et al*., 2018). Validation metrics to assess the model quality were computed with the Phenix comprehensive validation (cryo-EM) algorithm.

After generating the whole AU structure, Chimera *findclash* (integrated in the Scipion protocol *chimera-contacts*) was executed to identify possible interactions between pairs of chains contained in the AU or between a chain in the AU and a chain from a neighboring AU. The default parameters *(cutoff* = −0.4 and *allowance* = 0.0) were used to report as possible bonds all pair of atoms separated by a distance no higher than the sum of the corresponding van der Waals radii plus 0.4 Å. To identify additional unmodeled densities, we generated remnant maps by masking off from the initial map a region within a 3, 4 or 8 Å radius from the modelled atoms using ChimeraX (*vol zone invert true*). Cα backbones of the unmodeled densities were built by automatic placing of poly-A helices in Coot. Assignment of sequence of probable candidates was semi-automatically performed with the docking GUI in Coot and manually assessed.

Protein charge at pH 7.0 was calculated as described in http://isoelectric.org/index.html, with amino acid pKa values taken from (Lide and Haynes, 2009), and general pKa values for terminal amino and carboxy groups taken from (Berg *et al*., 2015). Protein sequence alignments were performed with Clustal Ω (http://www.clustal.org/omega/) integrated in Geneious version 2019.0 (https://www.geneious.com). Secondary structure elements and similarity depictions were performed with ESPript 3 (https://espript.ibcp.fr/ESPript/cgi-bin/ESPript.cgi).

Molecular graphics and analyses including molecule superposition (*matchmaker* command), all-atom RMSD calculation, map rendering, map normalization and colouring by surface electrostatic potential were performed with UCSF Chimera or UCSF ChimeraX. UCSF Chimera *Hide dust* was used for clarity when composing figures representing density maps.

## Data availability

The FAdV-C4 cryo-EM maps have been deposited at the Electron Microscopy Data Bank (EMDB, http://www.ebi.ac.uk/pdbe/emdb) with accession numbers EMD-19396 (AG234 strain, pentonless) and EMD-19401 (KR5 strain, full virion). The FAdV-C4 molecular model is deposited at the Protein Data Bank (PDB, https://www.ebi.ac.uk/pdbe/) with accession number 8ROQ.

## Acknowledgements

Work supported by grants from the Spanish State Research Agency (AEI/10.13039/501100011033), with co-funding from the European Regional Development Fund (BFU2013-41249-P, BFU2016-74868-P, PID2019-104098GB-I00 and PID2022-136456NB-I00) to C.S.M., and the Christian Doppler Research Association (grant no: 189) to IPOV. C.S.M. further acknowledges support by the European Innovation Council (Horizon Europe) under grant agreement No 101098647 (project iAds), and Marie Skłodowska-Curie Actions (grant agreement 101129778, project INVECTA). The C.S.M. group is a member of the Spanish Adenovirus Network (RED2022-134221-T/AEI/10.13039/501100011033), CSIC LifeHub, and CSIC BCBHub. The CNB-CSIC was further supported by AEI Severo Ochoa Excellence grants SEV-2013-0347, SEV-2017-0712 and CEX2023-001386-S). M.P.-I. was supported by a predoctoral contract from La Caixa Foundation (ID 100010434), under agreement LCF/BQ/SO16/52270032), and by the VIRMAT project from the Madrid Regional Government and the REACT-EU program. M.H.-P. holds a Ramón y Cajal position (RyC2021-030929-I) funded by MCIN/AEI/10.13039/501100011033 and European Union NextGeneration EU/PRTR. M.H.-P. also acknowledges grant PID2023-151078OB-I00 funded by MCIN/AEI/10.13039/501100011033 and the European Union NextGenerationEU/PRTR.

We thank the CNB-CSIC Proteomics, Electron and Cryo-electron Microscopy facilities (particularly F.J. Chichón) for excellent technical support; as well as Diamond Light Source for Titan Krios data collection at the UK national electron Bio-Imaging Center (eBIC) under proposal EM15997. We are also grateful to Ana Marek for helpful discussions at the beginning of this project.

## Author contributions

C.S.M. and M.H. designed research. M.P.-I., A.S., M.M., G.N.C, M.H.-P., R.M., M.H. and C.S.M. performed research. M.P.-I., M.M., M.H.-P., A.P., R.M. and C.S.M. analysed data. M.P.-I. and C.S.M. wrote the paper, with input from all other authors.

## Footnotes

1 Fowl adenovirus species C has recently been renamed as *Aviadenovirus hydropericardii*, according to the latest ICTV name system (https://ictv.global/report/chapter/adenoviridae/adenoviridae/aviadenovirus).

