## Supplementary material for "Aviadenovirus structure: a highly thermostable capsid in the absence of stabilizing proteins"

[3] Escuela Politécnica Superior. Universidad Autónoma de Madrid. 28049 Madrid (Spain)

[4] Clinic for Poultry and Fish Medicine, Department for Farm Animals and Food System Science, University of Veterinary Medicine, Vienna, Austria

[5] Present address: Department of Virology, Aggeu Magalhaes Institute. Oswaldo Cruz Foundation- (FIOCRUZ). PE-50740-465 Recife (Brazil)

[6] Present address: Institute of Medical Genetics, Center for Pathobiochemistry and Genetics, Medical University of Vienna. Währingerstrasse 10, 1090 Vienna, Austria

[7] Present addresses: Department of Materials Physics, Instituto de Ciencia de Materiales Nicolás Cabrera, and Condensed Matter Physics Center (IFIMAC). Universidad Autónoma de Madrid, Ciudad Universitaria de Cantoblanco, 28049, Madrid, Spain.

**Short title: Structure of an aviadenovirus, FAdV-C4**

**This PDF file includes:**

Supplementary text

Supplementary tables S1 to S18

Supplementary figures S1 to S9

Legend for supplementary file S1

Supplementary references

### Supplementary Text

*Structure of FAdV-C4 strain AG243 lacking pentons*

In the process of obtaining a high resolution map of FAdV-C4, a collection of cryo-EM grids prepared under different conditions was screened to select the grid that would contain the optimal ice thickness, particle concentration and intact morphology. One of the grids, prepared from samples of FAdV-C4 strain AG243, was selected and a 3.2 Å resolution map was obtained (**Table S2, Figure S2a-b**). Unfortunately, although visual inspection of the micrographs did not indicate that the viral particles were damaged, this map showed a capsid completely devoid of pentons (**Figure S2c**). It is known that adenovirus capsid disassembly starts at the pentons (Greber *et al.*, 1993; Ortega-Esteban *et al.*, 2013) and it is relatively common for adenovirus cryo-EM maps to present a low occupancy of pentons (Cheng *et al.*, 2014; Marabini *et al.*, 2021; Marsh *et al.*, 2006).

Prior works reported that upon freeze-thawing (Martinez *et al.*, 2015), heating or acidification (Pérez-Berná *et al.*, 2012), loss of peripheral core materials (presumably protein VI or core protein V) accompanied the release of pentons in HAdVC-5. Apart from the density corresponding to the pentons, our FAdV-C4 pentonless map is also lacking density accounting for connections between capsid and core (**Figure S2c, arrowheads**). These connections arise from the inner hexon cavity, where protein VI is located (Dai *et al.*, 2017; Hernando-Pérez *et al.*, 2020; Yu *et al.*, 2022). The loss of internal components at the outermost part of the core is also evidenced by a large dip in the radial average profile of the maps (**Figure S2d, arrowheads**). Another intriguing difference is the loss of densities near the 2-fold symmetry axes (designated as *RD4* in the main text, **Figure 5**) in the pentonless map (**Figure S2c, dots**). The identity and role of this RD is unknown.

*Aviadenovirus fibres*

Due to their flexibility and symmetry mismatch with the penton base, fibres cannot be resolved by cryo-EM single particle averaging when imposing icosahedral symmetry as we have done here, and thus a single blurry stump can be observed, resulting from averaging the proximal end of the two fibres located in each penton vertex (**Figure S2c left, pentagon**).

Since aviadenoviruses have two trimeric fibres per vertex (Benkö *et al.*, 2022; Gelderblom and Maichle-Lauppe, 1982; Hess *et al.*, 1995), their interaction with the penton base is expected to be different from other AdVs. A FNPVYPY sequence motif at the fibre N-terminal region has been shown to be involved in attachment to penton base and is conserved in mastadenoviruses (Zubieta *et al.*, 2005). The N-terminal sequences of both FAdV-C4 fibres differ from those in the HAdVs (**Figure S4b, bottom**). While in HAdV-C5 and HAdV-D26 the penton-binding conserved motif (FNPVYPY) is located very close to the protein N-terminus (starting approximately at residue 10), in both FAdV-C4 fibres the equivalent penton-binding peptide (LDLVYPF) is located further downstream, starting only at residue 70 in the short fibre and 60 in the long one (**Figure S4b, dashed rectangle**). A long poly-Gly stretch in the FAdV-C4 short fibre (**Figure S4b, black rectangle**) has been proposed to provide increased flexibility at the region between the N-terminal peptide and the start of the shaft (Zubieta *et al.*, 2005).

The N-terminal tails of fibres contact the penton base at the groove between monomers (Zubieta *et al.*, 2005). In our FAdV-C4 map, remnant density on top of the penton base shows a "starfish" shape corresponding to the 5-fold averaged fibre N-termini, similar to density observed in other adenovirus structures. A cylindrical density protruding outwards and corresponding to the 5-fold averaged start of the shafts is also observed (**Figure S4c**). Ten of the eighteen N-terminal amino acids of the HAdV-D26 fibre (containing the FNPVYPY motif) can be docked into the FAdV-C4 remnant density (**Figure S4c**). No additional density that could account for the longer N-terminal tails of the FAdV-C4 fibres (**Figure S4c**), or for the second fibre binding to a different location on the pentamer surface, was observed. In HAdV-D26, it has been proposed that the fibre N-terminal peptide (Ala2-Ala20) folds around the HVL at the periphery of the penton base pentamer **(Figure S4d, left**) (Yu *et al.*, 2017). If we overlap the HAdV-D26 fibre N-terminal peptide with the FAdV-C4 penton base structure, we observe that the HVL, VL and VL' loops, which fold back towards the main body of the pentamer instead of spreading out into the solvent, could provide additional interactions with the double fibre N-terminal tails and contribute to clasp them to the capsid (**Figure S4d, right**). Alternatively, this conformational difference could allow the accommodation of the extra-long N-terminal region preceding the penton binding motif (**Figure S4c**). Further studies using symmetry relaxation and localized reconstruction would be needed to solve the mode of binding of the two trimeric fibres to the pentameric pentons.

### Supplementary Tables

**Table S1.** FAdV-C4 proteins identified by LC-MS/MS

| Protein | Mass of immature protein (Da) | MASCOT Score | Total number of peptide (number of significant) matches | Sequences | Sum of peptide area (x$10^6$) |
|---|---|---|---|---|---|
| hexon | 106267 | 2203 | 52 (52) | 40 (40) | 86.41 |
| pX (pre-μ)[a] | 18649 | 1006 | 27 (27) | 22 (22) | 61.07 |
| pVIII[a] | 26956 | 749 | 18 (18) | 16 (16) | 8.69 |
| pIIIa[a] | 65275 | 649 | 16 (16) | 16 (16) | 2.91 |
| penton base | 57883 | 562 | 13 (13) | 13 (13) | 10.41 |
| pVI[a] | 24302 | 428 | 12 (12) | 9 (9) | 1.41 |
| pVII[a] | 9031 | 417 | 16 (16) | 12 (12) | 10.54 |
| L1 52/55k[a] | 44532 | 244 | 9 (9) | 8 (8) | 1.09 |
| fibre-1 | 45089 | 238 | 5 (5) | 5 (5) | 0.98 |
| fibre-2 | 50066 | 96 | 3 (3) | 3 (3) | 0.78 |
| IVa2 | 45810 | 56 | 2 (2) | 2 (2) | 0.09 |
| AVP | 24309 | 55 | 2 (2) | 2 (2) | 0.87 |
| pTP[a] | 70648 | 35 | 1 (1) | 1 (1) | 0.12 |

*[a]* Proteins cleaved by AVP during maturation (see **Figure S1**)

**Table S2.** Cryo-EM data collection, image processing, reconstruction and refinement for the two FAdV-C4 specimens used

| **Data collection** | **FAdV-C4 KR5** | **FAdV-C4 AG234** |
|---|---|---|
| Microscope | Titan Krios | Titan Krios |
| Camera | Falcon III (linear) | Falcon III (linear) |
| Voltage | 300 kV | 300 kV |
| Magnification | 97,902 | 95,890 |
| Nominal pixel size | 1.43 Å/px | 1.46 Å/px |
| Dose rate | 35 $e^{-}$/$Å^{2}$.s | 41$e^{-}$/$Å^{2}$.s |
| Exposure time | 1.1 s | 0.99 s |
| Cumulative electron dose | 38.6 e-/$Å^{2}$ | 40.5 e-/$Å^{2}$ |
| Number of frames | 39 | 39 |
| Defocus range | -1 to -2.5 µm | -1 to -2.5 µm |
| Micrographs collected | 4,306 | 4,796 |
| Acquisition software | EPU | EPU |
| **Image processing** | | |
| Frame alignment software | Motioncor2 Scipion | Motioncor2, Scipion |
| CTF estimation software | gCTF, Scipion | gCTF, Scipion |
| Particle picking software | XMIPP, Scipion | XMIPP, Scipion |
| Micrographs used | 3,769 | 4,733 |
| Particles picked | 13,587 | 40,383 |
| Particles after screening | 10,919 | 40,345 |
| Particles after 2D classes | 10,877 | 40,169 |
| **Reconstruction** | | |
| Software | RELION, Scipion | RELION, Scipion |
| Particles included | 9,466 | 32,425 |
| Symmetry imposed | Icosahedral | Icosahedral |
| Rotational accuracy | 0.05 degrees | 0.05 degrees |
| Translational accuracy | 0.11 pixels | 0.1 pixels |
| B-factor applied | -196 $Å^2$ | -211 $Å^2$ |
| Final resolution (gold standard FSC=0.143) | 3.3 $Å^2$ | 3.2 $Å^2$ |
| Experimental pixel size | 1.375 Å/px | 1.375 Å/px |

**Table S3.** Modelling and validation statistics for FAdV-C4 KR5 [a]

| **Model building, refinement and validation** | |
|---|---|
| Software | Scipion, Xmipp, UCSF Chimera, UCSF ChimeraX, Coot, Phenix |
| Chains | 16 |
| Atoms | 97638 (Hydrogens: 0) |
| Residues | Protein: 12237 Nucleotide: 0 |
| Water | **0** |
| Ligands | **0** |
| **Bonds (RMSD); outliers >4σ** | |
| Length (Å) | 0.005; 0 |
| Angles (°) | 1.03; 8 |
| MolProbity score | 2.21 |
| Clash score | 16.6 |
| **Ramachandran plot (%)** | |
| Outliers | 0.04 |
| Allowed | 8.02 |
| Favored | 91.94 |
| **Rama-Z (Ramachandran plot Z-score, RMSD)** | |
| whole | (N= 12201) -2.02 (0.07) |
| helix | (N= 1835) -1.18 (0.11) |
| sheet | (N= 2563) -1.05 (0.11) |
| loop | (N= 780.3) -1.53 (0.07) |
| **Rotamer outliers (%)** | 0.04 |
| **Cβ outliers (%)** | 0 |
| **Peptide plane** | |
| Cis proline/general | 0.0/0.0 |
| Twisted proline/general | 0.3/0.0 |
| **CaBLAM outliers (%)** | 5.33 |
| **ADP (B-factors)** | |
| Iso/Aniso | 97638/0 |
| min/max/mean | |
| Protein | 24.96/94.00/41.69 |
| **Model vs. Data** | |
| CC (mask) | 0.85 |
| CC (box) | 0.49 |
| CC (peaks) | 0.29 |
| CC (volume) | 0.81 |

[a]Obtained from Phenix v19 cryo-EM comprehensive validation. No model was built for FAdV-C4 AG234

**Table S4.** FAdV-C4 proteins traced in the model

| Protein | Length (amino acids) | Copy number in AU | Chain ID | Residues traced | Not traced | Number traced |
|---|---|---|---|---|---|---|
| **hexon** | 937 | 12 | A | 11-937 | 1-10 | 927 |
| | | | B | 2-935 | 1; 936-937 | 934 |
| | | | C | 4-937 | 1-3 | 934 |
| | | | D | 10-937 | 1-9 | 928 |
| | | | E | 11-937 | 1-10 | 927 |
| | | | F | 2-936 | 1; 937 | 935 |
| | | | G | 8-937 | 1-7 | 930 |
| | | | H | 11-935 | 1-10; 936-937 | 925 |
| | | | I | 11-937 | 1-10 | 927 |
| | | | J | 11-936 | 1-10; 937 | 926 |
| | | | K | 2-937 | 1 | 936 |
| | | | L | 11-937 | 1-10 | 927 |
| **penton base** | 525 | 1 | M | 54-524 | 1-53; 525 | 471 |
| **IIIa** | 590 | 1 | N | 16-259 | 1-16; 259-590 | 244 |
| **IIIa APD*** | | | | 305-384 | | 80 |
| **VIII** | 247 | 2 | O | 1-115; 174-241 | 116-173; 242-247 | 183 |
| | | | P | 1-115; 174-241 | 116-173; 242-247 | 183 |

*Tentative assignment (**Figure S6b**).

**Table S5.** Comparison between HAdV-C5 and FAdV-C4 virion proteins

| | Length (aa) | | Molecular weight (kDa) | | Isoelectric Point | | Charge at pH 7 | | % sequence homology | |
|---|---|---|---|---|---|---|---|---|---|---|
| | HAdV-C5 | FAdV-C4 | HAdV-C5 | FAdV-C4 | HAdV-C5 | FAdV-C4 | HAdV-C5 | FAdV-C4 | Pairwise identity | Pairwise Positive |
| **hexon**[a] | 952 | 937 | 108.007 | 106.049 | 4.94 | 4.88 | -23.94 | -19.22 | 44.5 | 58.1 |
| **penton base**[a] | 571 | 525 | 63.293 | 57.406 | 5.15 | 4.72 | -12.33 | -10.29 | 37.9 | 50.5 |
| **IIIa**[a] | 585 | 590 | 65.253 | 65.259 | 5.71 | 6.66 | -7.05 | -0.64 | 24.4 | 37.7 |
| **VIII**[a] | 227 | 247 | 24.687 | 26.859 | 9.24 | 6.15 | 2.41 | -1.02 | 22.5 | 36.8 |
| **VI**[b] | 250 | 227 | 26.996 | 24.26 | 10.42 | 11.02 | 8.06 | 11.35 | 19 | 34.4 |
| **VII**[b] | 198 | 77 | 21.992 | 9.037 | 12.45 | 13.48 | 40.31 | 24.91 | 14.5 | 19 |
| **V** | 368 | absent | 41.447 | absent | 10.75 | absent | 27.34 | absent | -- | -- |
| **μ** | 80 | 179 | 8.846 | 18.66 | 12.98 | 12.21 | 17.43 | 27.91 | 14.8 | 18.6 |
| **52/55k** | 415 | 399 | 47.06 | 44.389 | 5.47 | 5.08 | -10.01 | -10.47 | 24 | 37.6 |
| **fibre 1** | 581 | 433 | 61.585 | 45.059 | 6.01 | 4.53 | -3.51 | -7.95 | -- | -- |
| **fibre 2** | absent | 479 | absent | 49.812 | absent | 4.07 | absent | -20.01 | -- | -- |
| **AVP** | 204 | 209 | 23.068 | 24.039 | 8.21 | 9.29 | 5.17 | 7.27 | 36.7 | 63.8 |
| **TP** | 671 | 602 | 76.5 | 70.407 | 5.91 | 6.75 | -7.3 | -1.09 | 27.9 | 43.6 |
| **IVa2** | 449 | 394 | 50.887 | 45.383 | 8.69 | 7.41 | 8.34 | 1.58 | 28 | 43.3 |
| **IX**[b] | 140 | absent | 14.458 | absent | 6.45 | absent | -0.09 | absent | -- | -- |

[a] Proteins traced (totally or in part) in both HAdV-C5 and FAdV-C4

[b] Proteins traced only in HAdV-C5 model (Dai *et al.*, 2017)

AVP: adenovirus protease. TP: Terminal protein. Hexon, penton base, VII, 52/55k, TP and IVa2 are shorter in FAdV-C4 compared to HAdV-C5, whereas IIIa, VIII, μ and AVP are longer. The FAdV-C4 virion has two shorter and more acidic fibres compared to the single fibre in HAdV-C5. Proteins IX and V contribute outer negative and inner positive charges in HAdV-C5, but they are not present in the FAdV-C4 virions.

**Table S6.** Regions of hexon with differences between HAdV-C5 and FAdV-C4[a].

| Different region[b] | Amino acids in HAdV-C5 | Amino acids in FAdV-C4 | Observation |
|---|---|---|---|
| *diff 1* | absent | M1-T5 | N-term |
| *diff 2* | W135-V168 | S140-A150 | HVR 1 |
| *diff 3* | G176-K198 | Y158-S176 | HVR 2[c] |
| *diff 4* | Y212-A220 | P190-L202 | HVR 3 |
| *diff 5* | K226-W227 | K207 | Surface exposed |
| *diff 6* | I249-Q261 | L229-T230 | HVR 4 |
| *diff 7* | S268-V284 | M237-G246 | HVR 5 |
| *diff 8* | P304-G316 | P267-N275 | HVR 6[c] |
| *diff 9* | G419-TR453 | A379-Y417 | HVR 7 |
| *diff 10* | N491-K493 | D457-T459 | Big insertion neighborhood |
| *diff 11* | ~V749 | V714-A715 | Pincer |
| *diff 12* | K810-Y811 | A776 | Pincer |
| *diff 13* | Y787 | Y756 | Inside hexon cavity |
| *diff 14* | ~L820 | S785-S798 | Big insertion |
| *diff 15* | ~W814 | S813-V816 | Big insertion neighborhood |
| *diff 16* | D857 | Q840-G842 | Pincer |
| *diff 17* | T952 | absent | C-term |

[a] Two criterions were used to determine these regions: residues exceeding an RMSD cutoff of 5 Å, or insertions/deletions in either of the two sequences.

[b] Different regions sorted by sequence order.

[c] *diff3* and *diff8* extend beyond the previously reported boundaries of HVR2 and HVR6 in HAdV-C5 (Rux *et al.*, 2003).

**Table S7.** Intra-monomer interactions established by hexon *big insertion*

| Amino acids in *big insertion* | Interacting amino acids |
|---|---|
| Ser785 | Gln799 |
| Tyr786 | Tyr449, Leu475, Val478, Gln493 |
| Pro788 | Arg448, Lys450, Leu475 |
| Asn792 | Ser454, Phe456 |
| Ser793 | Ser454 |
| Gly794 | Pro268, Ser452, Ser454, Pro815 |
| Glu795 | Pro268, Ser813, Trp814 |
| Gln796 | Phe451, Leu475, Gln799, Ser813 |
| Pro797 | Ser813 |

**Table S8.** Different regions of penton base in HAdV-C5 and FAdV-C4.

| Different region[a] | Amino acids in HAdV-C5 | Amino acids in FAdV-C4 | Observation |
|---|---|---|---|
| *diff 1* | M1-L36 | M1-M53 | At the start of the N-terminal arm, non-traced[b] |
| *diff 2* | S76-S81 | T93-D97 | Located at the base, interaction with hexon. Specificity in hexon-penton interaction? |
| *diff 3* | P153-Q158 | P165-V181 | VL: longer in FAdV-C4 |
| *diff 4* | A294-D397 | N318-D328 | HVL: Shorter and lacks RGD in FAdV-C4. If located as in HAdV-C5, it would clash against the VL |
| *diff 5* | ~D397 | D347-K352 | Insertion in FAdV-C4. Given its length, proposed to be another variable loop (VL') |
| *diff 6* | G413-Q416 | P367 | Surface exposed |
| *diff 7* | S458 | N407 | Located at the base, interaction with hexon. Specificity in hexon-penton interactions? |
| *diff 8* | T493 | T446 | Fibre rearrangement region |

[a]Different regions sorted by sequence order.

[b]*diff1* is defined by sequence comparison and not by the RMSD> 5 Å criterion

**Table S9.** Comparison of intra-monomer interactions established by penton base variable loop (VL) in HAdV-C5 and FAdV-C4

| **HAdV-C5** | | **FAdV-C4** | |
|---|---|---|---|
| **Amino acids in VL** | **Interacting amino acids** | **Amino acids in VL** | **Interacting amino acids** |
| Pro153<br>Gln158 | Leu152, Val159<br>Val 159 | Pro165 | Asp 164, Asn223 |
| | | Pro166 | Arg163, Gln225 |
| | | Pro171 | His335, Gly339 |
| | | Pro172 | Leu334, Ser341 |
| | | Ser173 | Leu334 |
| | | Val175 | Pro332, Leu334,Val344,Tyr34, Pro353 |
| | | Gly176 | Val344, Ile345 |
| | | Tyr179 | Leu266, Pro267, Asn343, Ile345 |
| | | Val181 | Met160, Arg163, Gly183, Ala184 |

**Table S10.** Regions of protein VIII with RMSD> 5 Å between HAdV-C5 and FAdV-C4.

| **Different region[a]** | **Amino acids in HAdV-C5** | **Amino acids in FAdV-C4** | **Observations** |
|---|---|---|---|
| *diff 1* | S2-G4 | M1-A6 | Longer N-terminal in FAdV-C4 (interactions) |
| *diff 2* | A59-N103 | W60-V107 | Boundaries of α2' and α2'' in FAdV-C4 |
| *diff 3* | G158-Q172 | P174-V187 | β4 in HAdV-C5/ α2''' in FAdV-C4 (neck) |
| *diff4* | ~S222 | V233-G236 | Insertion in FAdV-C4 near C-terminus (interactions) |
| *diff 5* | Y226-D227 | E242-G247 | Flexible and non-traced in FAdV-C4 (interactions) |

[a]Different regions sorted by sequence order.

**Table S11.** Different regions of protein IIIa in HAdV-C5 and FAdV-C4.

| Different region[a] | Amino acids in HAdV-C5 | Amino acids in FAdV-C4 | Observation |
|---|---|---|---|
| *diff 1* | D4-N41 | A 19-A31 | Shorter in FAdV-C4 and flexible |
| *diff 2* | R128-G135 | V 117-R 126 | End of the connecting helix and loop to VIII-binding domain, causing its drastic orientation change. FAdV-C4 has a 2 amino acid insertion |
| *diff 3*[b] | R193-Q194 | M184-G186 | Handle |
| *diff 4*[b] | A 216-S225 | W208-G217 | Not traced in HAdV-C5 but forms alpha helix in FAdV-C4 |
| *diff 5*[b] | A 268-L301 | … | Not ordered in FAdV-C4 |

[a]Different regions sorted by sequence order.
[b]*diff 3, 4 and 5* were defined by comparison of the structures after superposition of the VIII-binding domains

**Table S12.** Interactions between hexons in the same facet (ST interfaces). For interface nomenclature, see **Fig. S9.**

| ST1: H2—H1 | | | |
|---|---|---|---|
| E | Ile76 | Ile76 | A |
| | Gln77 | Tyr74, Ile76 | |
| | Asp79 | Arg72, Tyr74 | |
| | Arg86 | Tyr74 | |
| | Glu311 | Asn92, Arg919 | |
| | Arg312 | Gly94,Asp95,Arg919 | |
| | Ser313 | Asp95 | |
| | Gly314 | Gly94, Trp97 | |
| | Met315 | Lys65-Gln67,Trp97,His589 | |
| | Glu547 | Tyr74 | |
| | Ala936 | Glu64 | |
| | Val937 | Thr62, Glu64 | |
| E | Thr81-Thr83 | Thr718 | C |
| | Arg86 | Glu719 | |
| | Gly302 | Pro705 | |
| | Val303 | Leu703, Thr704,Phe721 | |
| | Val319 | Asp700 | |
| | Pro322 | Asn706 | |
| | Lys615 | Ser693 | |
| | Pro633 | Ser886 | |
| | Ala634 | Ala634 | |
| | Arg635 | Asp691, Thr692 | |
| | Gln659 | Glu719 | |
| | Val662 | Leu716 | |
| | Arg928 | Ser693 | |
| | Phe931 | Thr692,Ser693 | |
| | Ala932 | Ser693,Ile694 | |
| | Gly934 | Ile694,Met879 | |
| | Asn935 | Asn695-Asn699,Tyr880 | |
| | Ala936 | Asn877,Met879 | |
| | Val937 | Asn699, Arg701,Ser863 | |
| F* | Leu4 | Asn60,Gln594 | A* |

| ST2: H1—H4 | | | |
|---|---|---|---|
| A | Arg88 | Tyr74,Pro75,Glu577 | L |
| | Arg312 | Asn92,Gly94 | |
| | Gly314 | Ile71 | |
| | Met315 | Arg68 | |
| | Glu547 | Tyr74 | |
| A | Arg86 | Thr718 | K |
| | Val303 | Leu716,Thr718 | |
| | Cys304 | Thr718 | |
| | Met315 | Asp700 | |
| | Val319 | Pro705 | |
| | Pro633 | Thr629 | |
| | Ala634 | Asn631 | |
| | Thr636 | Thr692 | |
| | Glu638 | Thr692 | |
| | Gln659 | Leu716,Thr718 | |
| | Ala932 | Ser693 | |
| | Thr933 | Ser687,Leu689,Asn890 | |
| | Asn935 | Asn890 | |
| B* | Thr9 | Gly698 | |
| | Arg14 | Ile694 | |
| | Asp7 | Lys65,Ala66 | L* |

| ST3: H2—H3 | | | |
|---|---|---|---|
| | Ile76 | Ile76 | G |
| | Gln77 | Tyr74,Ile76 | |
| | Asp79 | Arg72,Tyr74 | |
| | Arg86 | Tyr74 | |
| | Arg312 | Gly94,Asp95,Arg919 | |
| | Gly314 | Gly94 | |
| | Met315 | Lys65-Gln67,Trp97,His589 | |
| | Glu547 | Tyr74 | |
| | Val937 | Thr63,Glu64 | |
| D | Ser82 | Thr718 | I |
| | Thr83 | Thr718 | |
| | Arg86 | Glu719 | |
| | Gly302 | Pro705 | |
| | Val303 | Leu703-Pro705 | |
| | Val319 | Asp700 | |
| | Pro322 | Asn706 | |
| | Lys615 | Thr692,Ser693 | |
| | Ile632 | Thr692 | |
| | Pro633 | Thr692,Ser886 | |
| | Arg635 | Asp691-Ser693 | |
| | Gln659 | Glu719 | |
| | Arg928 | Ser693 | |
| | Phe931 | Thr692,Ser693 | |
| | Ala932 | Ser693,Ile694 | |
| | Asn935 | Ile694-Trp696,Met879,Tyr880 | |
| | Ala936 | Asn877,Met879 | |
| | Val937 | Asn699,Ser863,Leu874 | |

(table continues in next page)

**Table S12 (continued)**

**ST4: H4—H2**

| Chain | Residue | Interacting residues | Chain |
|---|---|---|---|
| L | Ile76 | Ile76 | E |
| | Gln77 | Tyr74,Ile76 | |
| | Thr78 | Tyr74 | |
| | Asp79 | Arg72,Tyr74 | |
| | Arg86 | Tyr74 | |
| | Arg88 | Tyr74 | |
| | Glu311 | Arg919 | |
| | Arg312 | Gly94,Asp95,Arg919 | |
| | Gly314 | Gly94,Trp97 | |
| | Met315 | Lys65-Trp97,His589 | |
| | Val319 | Ala66 | |
| | Val937 | Thr62,Glu64 | |
| | Ser82 | Thr718 | D |
| | Thr83 | Thr718 | |
| | Gly302 | Pro705 | |
| | Val303 | Thr704,Pro705 | |
| | Val319 | Asp700 | |
| | Pro322 | Asn706 | |
| | Lys615 | Thr692 | |
| | Ile632 | Thr692 | |
| | Pro633 | Thr692,Ser886 | |
| | Arg635 | Asp691,Thr692 | |
| | Gln659 | Glu719 | |
| | Arg928 | Ser693 | |
| | Phe931 | Thr692,Ser693 | |
| | Ala932 | Ser693,Ile694 | |
| | Thr933 | Ser693,Ile694 | |
| | Asn935 | Ile694-Gly698 | |
| | Ala936 | Leu874,Asn877,Met879,Tyr880 | |

**ST5: H3—H4**

| Chain | Residue | Interacting residues | Chain |
|---|---|---|---|
| I | Ile76 | Ile76 | J |
| | Gln77 | Tyr74,Ile76 | |
| | Thr78 | Tyr74 | |
| | Asp79 | Arg72,Tyr74 | |
| | Arg86 | Tyr74 | |
| | Arg312 | Gly94,Asp95,Arg919 | |
| | Ser313 | Gly94 | |
| | Gly314 | Gly94,Trp97 | |
| | Met315 | Lys65,Ala66,Gln67,Trp97,His589 | |
| | Val319 | Ala66 | |
| | Glu547 | Tyr74 | |
| | Val937 | Glu64 | |
| | Ser82 | Thr718 | L |
| | Thr83 | Thr718 | |
| | Gly302 | Pro705,Asn706 | |
| | Val303 | Thr704,Pro705 | |
| | Val319 | Asp700 | |
| | Pro322 | Asn706 | |
| | Lys615 | Thr692,Ser693 | |
| | Ile632 | Thr692 | |
| | Pro633 | Thr692,Ser886 | |
| | Arg635 | Asp691,Thr692 | |
| | Gln659 | Glu719 | |
| | Arg928 | Ser693 | |
| | Phe931 | Thr692,Ser693 | |
| | Ala932 | Ile694 | |
| | Thr933 | Ser693,Ile694 | |
| | Gly934 | Met879 | |
| | Asn935 | Ile694,Trp696,Asn699,Met879,Tyr880 | |
| | Ala936 | Leu874,Asn877,Met879 | |
| | Val937 | Asn699,Ser863,Leu874,Asn877 | |

**ST6: H3—H3 (AU1)**

| Chain | Residue | Interacting residues | Chain |
|---|---|---|---|
| H | Ile76 | Ile76 | I’ |
| | Gln77 | Tyr74,Ile76 | |
| | Asp79 | Arg72,Tyr74 | |
| | Arg86 | Tyr74 | |
| | Glu311 | Arg919 | |
| | Arg312 | Asp95,Arg919 | |
| | Gly314 | Gly94,Trp97 | |
| | Met315 | Lys65 | |
| | Val319 | Ala66 | |
| | Asp691 | Arg635 | H’ |
| | Thr692 | Ile632,Pro633,Arg635,Phe931 | |
| | Ser693 | Lys615,Arg928,Phe931,Ala932,Thr933 | |
| | Ile694 | Ala932,Thr933,Gly934 | |
| | Asp700 | Val319 | |
| | Thr704 | Val303 | |
| | Pro705 | Gly302,Val303 | |
| | Asn706 | Gly302,Pro322 | |
| | Leu716 | Val662 | |
| | Thr718 | Thr81,Ser82 | |
| | Glu719 | Arg86,Gln659 | |
| | Asn877 | Asn935 | |
| | Met879 | Asn935 | |
| | Tyr880 | Asn935 | |
| | Ser886 | Pro633 | |

(table continues in next page)

**Table S12 (continued)**

| ST7: H4—H3 (AU 1) | | | |
|---|---|---|---|
| J | Arg88 | Tyr74 | H' |
| J | Glu311 | Asn92 | H' |
| J | Arg312 | Asn92 | H' |
| J | Met315 | Gln67,Arg68 | H' |
| J | Glu547 | Tyr74 | H' |
| J | Arg86 | Thr718 | G' |
| J | Val303 | Leu716,Thr718 | G' |
| J | Cys304 | Thr718 | G' |
| J | Met315 | Asp700 | G' |
| J | Val319 | Pro705 | G' |
| J | Pro633 | Thr629 | G' |
| J | Ala634 | Asn631 | G' |
| J | Thr636 | Thr692 | G' |
| J | Phe927 | Ser693 | G' |
| J | Ala932 | Ser693 | G' |
| J | Thr933 | Ser687,Leu689,Asn890 | G' |
| K* | Asp7 | Asn699 | G' |
| K* | Thr9 | Gly698,Asn699 | G' |
| K* | Thr10 | Asn699 | G' |
| K* | Asp7 | Ala66 | H'* |
| K* | Thr9 | Ala66 | H'* |

Interacting amino acids were identified with UCSF Chimera *findclash* (Pettersen *et al.*, 2004) implemented in Scipion (Martínez *et al.*, 2020). Following the nomenclature in (Liu *et al.*, 2010), **ST**, **TT** and **SS** indicate the three kinds of interfaces between hexons (**Fig. S9, see also Tables S13 and S14**). **S** refers to the facet of the hexon trimer pseudo-hexagonal base composed by the two β-barrels in a single monomer; **T** refers to the facet composed by two β-barrels coming from two different hexon monomers. **H1-H4** refer to the four hexon trimers in the icosahedral asymmetric unit. Suffixes _AU1 to _AU7 indicate neighbouring asymmetric units (**Fig. S9**). Letters **A-L** (blue shaded columns, see also **Fig. S9**) denote the 12 hexon monomer chains in the icosahedral asymmetric unit. A prime (') symbol indicates chains belonging to the neighbouring asymmetric units. **Purple text** indicates residues potentially involved in salt bridges and underlining indicates recurrent salt bridges. Cells shaded in light orange indicate C-terminal flexible regions and pink shading indicates N-terminal flexible regions. These flexible regions were defined based on the RMSD analysis of the twelve hexon chains of the asymmetric unit (**Fig. S3c**). Residues involving difference regions of FAdV-C4 and HAdV-C5 (**Table S6 and Fig. 2a**), except those at the N- or -C termini, where not found in the interaction analysis, indicating that interactions between hexons are conserved between FAdV-C4 and HAdV-C5. Only the N- and -C termini, intrinsically flexible and variable, establish different interactions.

*Notice that some "S" interfaces, which are defined as involving a single hexon monomer on the basis of the hexagonal shape of the trimer, in fact may involve residues from two different monomers. This is due to the extensive interlacing of molecules in the hexon trimer, which results in the N-terminus of one hexon monomer reaching all the way to the center of the hexagon facet formed by the adjacent monomer (Rux *et al.*, 2003).

**Table S13.** Interactions between hexons in different facets (TT interfaces). For interface nomenclature, see **Fig. S9.** Nomenclature and colour codes as in Table S12.

**Local two-fold axes**

| TT8: H1—H1 (AU 3) | | | |
|---|---|---|---|
| C | Glu64<br>Lys65<br>Ala66 | Lys65, Ala66<br>Glu64, Ala66<br>Ala66 | B’ |
| C | Lys65<br>Ala66<br>Arg68<br>Ile71<br>Asn90<br>Asn92<br>Asp95<br>Trp97<br>Arg312<br>Arg919 | Asn699<br>Gly698-Asp700<br>Asn706<br>Ser693<br>Thr692<br>Asp691,Thr692,Ser693<br>Met879,Asn883<br>Ile694<br>Ala634<br>Asn883-His885 | A’ |
| B | Gly698<br>Asn699<br>Asp700 | Asp700<br>Asp700<br>Asn706 | A’ |
| B | Asp691<br>Thr692<br>Ser693<br>Ile694<br>Met879<br>Asn883<br>Ser884<br>His885<br>Ser886 | Asn92<br>Arg72,Phe73,Tyr74<br>Arg72,Tyr74<br>Arg68,Ile71<br>Gly94,Asp95,Trp97<br>Asn92-Gly94,Ser541<br>Glu311,Ser543<br>Asn92<br>Asn90 | B’ |

| TT9: H2- H4 (AU 3) | | | |
|---|---|---|---|
| F | Glu64<br>Lys65<br>Ala66 | Lys65, Ala66<br>Glu64, Ala66<br>Ala66 | K’ |
| E | Asp691<br>Thr692<br>Ser693<br>Ile694<br>Asn699<br>Asn883<br>Ser884 | Asn92<br>Tyr74,Ile76<br>Arg72,Tyr74<br>Arg68<br>Ala66<br>Asp95, Arg919<br>Asn92, Glu311 | K’ |
| F | Lys65<br>Ala66<br>Ile71<br>Asn90<br>Asn92<br>Gly94<br>Asp95<br>Arg919 | Asn699<br>Gly698-Asp700<br>Ser693<br>Thr692<br>Asp691,Thr692<br>Ile694<br>Asn883, His885<br>Ser884,His885 | J’ |
| E | Gly698<br>Asn699<br>Asp700 | Asp700<br>Asp700<br>Pro705, Asn706 | J’ |

**Icosahedral 2-fold axis**

| TT10: H2—H2 (AU 5) | | | |
|---|---|---|---|
| D | Arg68<br>Ile71<br>Arg72<br>Phe73<br>Tyr74<br>Ile76 | Asn706<br>Ser693<br>Ser693<br>Thr692,Ser693<br>Leu689,Thr692,Asn890<br>Thr692 | F’ |
| F | Leu689<br>Thr692<br>Ser693<br>Asn706<br>Asn890 | Tyr74<br>Phe73,Tyr74,Ile76<br>Ile71,Arg72,Phe73<br>Arg68<br>Tyr74 | D’ |
| F | Ala627<br>Gly698<br>Asp700<br>Leu703<br>Pro705<br>Asn706<br>Thr718 | Thr718<br>Asp700<br>Gly698,Asp700<br>Asn706<br>Asn706<br>Leu703,Pro705<br>Ala627 | F’ |

Note that $TT_{10}$ has the lowest number of residues involved in interactions (28 residues), compared to $TT_8$ and $TT_9$ (with 43 and 33 residues interacting respectively). Moreover, $TT_{10}$ does not have any pair of residues that could chemically form a salt bridge, thus the $TT_{10}$ interaction is the weakest among the TT interactions. Interestingly, there is a remnant density (*RD4*) that could be strengthening the interaction (see text section “***Additional internal densities***”). The salt bridges at the local two-fold axes are conserved between genera (Marabini *et al.*, 2021)

**Table S14.** Interactions between hexons in different facets (SS interfaces). For interface nomenclature, see **Fig. S9.** Nomenclature and colour codes as in Tables S12.

Local two-fold axis

| **SS11: H4—H1 (AU 3)** | | | |
|---|---|---|---|
| C | Asp79 | Asn663 | K' |
| | Gly302 | Val303,Gln659 | |
| | Val303 | Thr83,Gln659,Tyr660 | |
| | Arg312 | Asn631,Ile632,Pro633 | |
| | Ser313 | Ile632 | |
| | Met315 | Asp323,Lys615 | |
| | Glu320 | Val303 | |
| | Ala602 | Ala936,Val937 | |
| | Thr603 | Val937 | |
| | Pro633 | Ile76,Gln77 | |
| | Arg635 | Arg88 | |
| | Thr636 | Gly314 | |
| | Gln659 | Val662 | |
| | Asn882 | Arg312 | |
| | Asn883 | Arg312 | |
| | Ser884 | Glu311 | |
| | Asn915 | Ala936 | |
| | Glu918 | Pro633,Ala634 | |
| | Asn920 | Pro633 | |
| | Val921 | Pro633 | |
| | Thr933 | Thr307 | |
| | Asn935 | Asp79 | |
| | Ala936 | Arg86,Glu547 | |
| | Val937 | Gln77 | |
| A* | Arg14 | Met315 | |

| **SS12: H2—H3 (AU 5)** | | | |
|---|---|---|---|
| F | Asp79 | Asn663 | G' |
| | Val303 | Tyr660,Val662 | |
| | Arg312 | Ile630,Asn631,Ile632,Pro633 | |
| | Met315 | Asp323,Lys615 | |
| | Val319 | Val303 | |
| | Glu320 | Val303 | |
| | Leu321 | Val303 | |
| | Ala602 | Ala936,Val937 | |
| | Asn631 | Gln77 | |
| | Pro633 | Ile76,Gln77,Arg88 | |
| | Ala634 | Arg88,Glu311 | |
| | Thr636 | Gly314 | |
| | Glu638 | Met315 | |
| | Gln659 | Val662 | |
| | Asn882 | Glu311,Arg312 | |
| | Asn883 | Arg312 | |
| | Ser884 | Glu311 | |
| | Asn915 | Ala936 | |
| | Glu918 | Ala634 | |
| | Asn920 | Pro633 | |
| | Val921 | Pro633 | |
| | Thr933 | Thr307 | |
| | Asn935 | Asp79,Arg86 | |
| | Ala936 | Thr81 | |
| D* | Arg14 | Met315 | |

*Notice that some "S" interfaces, which are defined as involving a single hexon monomer on the basis of the hexagonal shape of the trimer, in fact may involve residues from two different monomers. This is due to the extensive interlacing of molecules in the hexon trimer, which results in the N-terminus of one hexon monomer reaching all the way to the center of the hexagon facet formed by the adjacent monomer (Rux *et al.*, 2003).

**Table S15.** Interactions between hexon and penton base.

| H1-P | | | | P-H1(AU 3) | | | |
|---|---|---|---|---|---|---|---|
| B | Thr83 | Lys96 | M | M | | | B' |
| | Arg86 | Thr408 | | | | | |
| | Val303 | Ile91,Arg104 | | | | | |
| | Ser305 | Thr408 | | | | | |
| | Thr307 | Thr408 | | | Asp71 | Arg312 | |
| | Gly314 | Glu405 | | | Tyr72 | Glu311,Arg312 | |
| | Met315 | Glu405,Lys485 | | | Thr73 | Arg312 | |
| | Asp323 | Thr120 | | | | | |
| | Glu547 | Thr408 | | | | | |
| | Pro633 | Asp112,Leu113 | | | | | |
| | Ala634 | Arg70 | | | | | C'* |
| | Arg635 | Arg70,Leu113 | | | | | |
| | Thr636 | Asp71 | | | | | |
| | Gln655 | Tyr94 | | | | | |
| | Ala658 | Tyr94 | | | | | |
| | Gln659 | Asn95,Lys96,Asp97 | | | | | |
| | Tyr660 | Lys96 | | | Arg56 | Thr9,Thr10 | |
| | Asp661 | Tyr94 | | | Met58 | Leu4,Thr5,Pro6 | |
| | Asn663 | Tyr94 | | | | | |
| | Ala932 | Glu121 | | | | | |
| | Thr933 | Thr120,Ser122 | | | | | |
| | Gly934 | Ser122 | | | | | |
| | Asn935 | Tyr69,Ser122,Gln124 | | | | | |
| C* | Leu8 | Asn127 | | | | | |
| | Thr9 | Asn127,Arg128 | | | | | |

Nomenclature and colour codes as in the previous tables. **P** indicates the penton base monomer (with chain id M in the coordinate file). All hexon-penton interactions involve "S" facets in the hexon pseudohexagonal base (**Fig. S9**). Residues in regions differing from HAdV-C5 (***diff2* in Table S8**) are shadowed in yellow. Notice that, although some penton residues involved in the interaction with hexon, are part of the difference regions, this is not the case for their partners in hexon. This observation indicates that hexon residues interacting with penton base are conserved, but penton interacting residues differ.

*Notice that some "S" interfaces, which are defined as involving a single hexon monomer on the basis of the hexagonal shape of the trimer, in fact may involve residues from two different monomers. This is due to the extensive interlacing of molecules in the hexon trimer, which results in the N-terminus of one hexon monomer reaching all the way to the center of the hexagon facet formed by the adjacent monomer.

**Table S16.** Interactions between protein IIIa and other capsid components. Nomenclature and colour codes as in the previous tables.

| **IIIa interactions with hexon** | | | | | | | | | |
|---|---|---|---|---|---|---|---|---|---|
| **IIIa-H1** | | | | | **IIIa' (AU3)-H1** | | | | |
| | *Domain* | | | | | *Domain* | | | |
| N | VIII binding | Asp209<br>Ala210<br>Val211 | Arg26<br>Glu27<br>Arg26,Glu31 | A | N | GOS- glue | Tyr76<br>Asp78<br>Arg90<br>Trp94 | Arg59<br>Arg59<br>Ala11,Pro13<br>Thr10,Ala11 | C |
| N | GOS-glue | Pro38<br>Tyr39<br>Ala40<br>Glu60<br>Val66<br>Lys67<br>Pro77<br>Asp78<br>Met80<br>Gly81<br>Ala82<br>His84<br>Ser85<br>Leu88<br>Asn89 | Gln876<br>Gln876,Pro878<br>Leu871,Gln876<br>Asn601<br>Leu597<br>Leu597<br>Asn590,Asn593<br>His589,Asn590,Asn593<br>Asn593,Gln594,Leu597<br>Asn593,Glu596,Leu597<br>Asp95,Arg919<br>Leu597<br>Glu918,Arg919<br>Pro917,Glu918<br>Arg312,Glu918 | B | | | | | |
| N | VIII-binding | Arg150<br>Tyr237<br>Gly238<br>Met240<br>Pro242 | Arg26<br>Arg26,Glu31,Gln34,Gln35<br>Arg26<br>Arg26<br>Arg26 | B | | | | | |
| N | Core-binding | Arg251<br>Lys254 | Gln35<br>Glu31 | B | | | | | |
| N | GOS-glue | Pro38<br>Ala40<br>Asn41<br>Ile44<br>Gln47<br>Thr48<br>Val51<br>Pro53<br>Lys54<br>Asp56 | Val55,Ala56<br>Ala21<br>Ala21,Gly22,Val55<br>Pro13,Leu15,Gln16<br>Pro13<br>Pro13,Gln16<br>Ala11<br>Ala11,Thr12<br>Thr10<br>Pro6 | C | | | | | |
| N | VIII-binding | Tyr142<br>Lys143<br>Thr144<br>Asp146 | Glu31<br>Glu31,Gln35<br>Glu31,Gln34,Gln35<br>Arg26 | C | | | | | |

(table continues in next page)

**Table S16 (continued)**

| IIIa interactions with penton base | | | | |
|---|---|---|---|---|
| **IIIa (AU3)-P** | | | | |
| | *Domain* | | | |
| N | GOS-glue | Lys54<br>Val55<br>Asp56<br>Gly57<br>Arg61<br>Tyr92<br>Thr93<br>Trp94 | Arg56<br>Gln55,Arg56<br>Arg56,Met58<br>Gln55<br>Gln55<br>Pro60,Thr61<br>Thr61,Gly62<br>Arg128 | P |
| | Conecting helix | Asn95<br>Gln100 | Met58,Pro60<br>Val57 | |

(table continues in next page)

**Table S16 (continued)**

| IIIa interactions with other IIIa molecules | | | | | |
|---|---|---|---|---|---|
| **IIIa-IIIa’(AU3)** | | | | | |
| | *Domain* | | | | *Domain* |
| N | GOS glue | Val18 | Leu69 | N’ | GOS glue & connecting helix |
| | | Ala19 | Asp109 | | |
| | | Ala21 | Gln72 | | |
| | | Leu22 | Leu69,Gln72,Leu106,Asp109,Val110,Gly113 | | |
| | | Ser23 | Asp109,His112,Gly113 | | |
| | | Ser24 | Gln72,Gly113,Lys116 | | |
| | | His25 | Gln72,Lys116,Val117 | | |
| | | Ala26 | Gln72,Val117 | | |
| | | Ala31 | Gln72,Gly73,Ala74 | | |
| | | Leu34 | Ala74 | | |
| | | Arg35 | Gly73,Ala74,Tyr76 | | |
| | | Tyr39 | Gln79,Ile83 | | |
| | | Arg42 | Ala74,Ile75,Gln79 | | |
| | | Leu43 | Asp86 | | |
| | | Leu46 | Leu69,Ile75,Ile83 | | |
| | | Gln47 | Asp86,Arg90 | | |
| | | Met50 | Val65,Leu87,Val91,Ser102,Ile103,Leu106 | | |
| | | Val51 | Arg90,Val99 | | |

(table continues in next page)

**Table S16 (continued)**

| **IIIa interactions with protein VIII (chain O)** | | | | | | | | | | | |
|---|---|---|---|---|---|---|---|---|---|---|---|
| **IIIa-VIII** | | | | | | **IIIa-VIII' (AU3)** | | | | | |
| | *Domain* | | | *Domain* | | | *Domain* | | | *Domain* | |
| N | GOS glue | Glu71<br>Gly73<br>Tyr76<br>Pro77 | Tyr234<br>Tyr234<br>Val233<br>Val233 | Body | O | N | GOS glue | Ala26<br>Asn27<br>Arg35<br>Tyr36 | Tyr234<br>Tyr234<br>Tyr234<br>Lys235 | Body | O |
| | VIII binding | Asp236<br>Tyr237 | Ser191,Gln192<br>Ser191 | Neck | | | | | | | |

**Table S17.** Interactions between protein VIII (chain O) and hexons in the peripentonal region. For interactions between VIII and IIIa, see **Table S16**. Nomenclature and colour codes as in the previous tables.

**VIII-H1**

| | *Domain* | | | |
|---|---|---|---|---|
| O | Body | Asn16<br>Val18<br>Thr19<br>Ala23<br>Asp88<br>Val89 | Glu638,Asn882<br>Glu638,Phe927<br>Thr881<br>Asn882<br>Glu918<br>Leu597 | A |
| | Neck | Ser92<br>Gly93<br>Pro94 | Leu597<br>Asp95,Asn593<br>Asn590,Asn593 | |
| | Head | Leu112<br>Ser113<br>Gly114<br>Glu175<br>Thr177 | Glu64,Lys65<br>Lys65,Ala66<br>Lys65<br>Asn590<br>Gln594 | |
| | Neck | Phe181<br>Lys182<br>Leu185<br>Arg186<br>Val187<br>Gln188<br>Gly189<br>Pro190 | Gln594<br>Leu597<br>Gln594,Met598, Asn601<br>Leu597,Arg600, Asn601,Pro917<br>Asn601<br>Asn601,Thr603<br>Thr603<br>Thr603 | |
| | Body | Ala228<br>Phe240<br>Glu241 | Asn882<br>Asn877,Pro878,Met879<br>Met879 | |
| | Body | Pro17<br>Val18<br>Gln73<br>Pro75<br>Tyr76<br>Ala77<br>Ile85 | Thr12,Pro13<br>Pro13,Leu15<br>Asp7<br>Leu4,Thr5<br>Ala3,Leu4,Thr5<br>Ala3,Thr5<br>Ala3 | B |
| | Neck | Tyr184<br>Leu185<br>Val187<br>Gln188<br>Gly189<br>Pro190<br>Ser191<br>Gln192<br>Glu196 | Asp32<br>Asp32<br>Ala2,Ala3<br>Ser30,Asp32<br>Ser30<br>Tyr28<br>Leu29,Glu31<br>Glu27<br>Pro23 | |
| | Body | Val199<br>Ser201<br>Gln202<br>Phe230<br>Phe240<br>Glu241 | Gln16<br>Thr12<br>Thr12<br>Arg59<br>Thr58,Arg59<br>Glu64 | |
| | Head | Gly106<br>Val107<br>Arg109<br>Gln111 | Met879,Asn883, His885<br>His885<br>Met879<br>Gly698,Asn699 | C |

**VIII-H2**

| | *Domain* | | | |
|---|---|---|---|---|
| O | Neck | Gly93<br>Pro95<br>Ser97<br>Ala98<br>Val99<br>Pro101 | Arg312<br>Arg312,Glu918,Val921<br>Pro917,Glu918<br>Asn915,Pro917<br>Ala602,Asn915<br>Ala925 | E |
| | Head | Gln108<br>Arg109<br>Val110<br>Gln111<br>Ser113<br>Gly114<br>Gly115 | Arg635,Thr636, Glu638,Phe927, Thr929,Ala932<br>Ala932,Gly934,Ala936<br>Tyr926,Phe927<br>Val319,Tyr926,Thr933<br>Asn316,Val317,Met924<br>Ala923,Met924<br>Ser313,Met315,Leu922 | |
| | Neck | Tyr100<br>Pro101 | Asp7<br>Arg14,Tyr17 | F |
| | Head | Asp103<br>Val110<br>Pro174<br>Glu175<br>Met176<br>Thr177<br>Pro178 | Arg14<br>Phe18<br>Pro6,Asp7<br>Leu4,Thr5,Pro6<br>Leu4,Thr5,Asp7<br>Ala3<br>Ala3,Thr5 | |
| | Neck | Phe181 | Ala2 | |

(table continues in next page)

**Table S17 (continued)**

| VIII-H4 | | | | |
|---|---|---|---|---|
| | *Domain* | | | |
| O | Body | Ala7 | Val937 | K |
| | | Pro8 | Val937 | |
| | | Val12 | Asn882,Ser884 | |
| | | Trp13 | Asn883,Ser884 | |
| | | Lys14 | Ser884 | |
| | | Pro17 | Asp691,Ile694 | |
| | | Val18 | Ser693 | |
| | | Gln26 | Ser884 | |
| | | Asn28 | Thr636 | |
| | | Tyr29 | Ala634,Arg635,Ala932 | |
| | | Gly30 | Thr636,Thr929, Pro930,Ala932 | |
| | | Ala31 | Thr636,Ala932 | |
| | | Thr32 | Phe927,Ala932 | |
| | | Ile33 | Ala925,Phe927, Gly934,Asn935 | |
| | | Asp34 | Ala925,Tyr926,Gly934 | |
| | | Trp35 | Asn935,Val937 | |
| | | Val36 | Asn935 | |
| | | Leu37 | Met315,Asn316 | |
| | | Pro38 | Met315,Asn316, Val317,Val319, Ala923,Met924 | |
| | | Gly39 | Met315,Ala923,Met924 | |
| | | Gly40 | Met315 | |
| | | Ser42 | Pro917,Ala923 | |
| | | Phe43 | Arg913,Asn915, Ala923-Ala925 | |
| | | Arg51 | Val937 | |
| | | Thr60 | Asn882 | |
| | | Phe67 | Asn877,Pro878 | |
| | | Gln202 | Met879 | |
| | | Met206 | Pro878,Met879 | |
| | | Pro211 | Asn882 | |
| | | Ala31 | Arg14 | L |
| | | Ile33 | Phe18 | |
| | | Phe43 | Tyr17 | |
| | | Ala46 | Tyr17 | |
| | | Ile50 | Arg14,Tyr17 | |
| | | Arg53 | Pro13 | |
| | | Phe67 | Arg59 | |
| | | Glu70 | Arg59,Asn60 | |
| | | Ser71 | Arg59,Glu64 | |
| | | Asp72 | Thr63,Asp588,Asn590 | |
| | | Gln73 | Lys65 | |
| | | His80 | Asp95,Asn593 | |
| | | Glu81 | Lys65 | |
| | | Ile84 | Asp95 | |

| VIII (AU3)-H1 | | | | |
|---|---|---|---|---|
| | *Domain* | | | |
| O | Body | Met1 | Pro13,Gln16 | A |
| | | Asn2 | Gln16 | |
| | | Leu3 | Pro13,Gln16,Tyr17 | |
| | | Leu4 | Pro23,Tyr28 | |
| | | Val36 | Pro13,Arg14,Tyr17 | |
| | | Leu37 | Phe18 | |
| | | Ala44 | Pro13 | |
| | | Phe218 | Ser30,Asp32 | |
| | | Asp219 | Glu31 | |
| | | Tyr11 | Asn601 | C |
| | | Lys14 | Glu918 | |
| | | Gln26 | Glu918 | |
| | | Gln27 | Asn601,Ala602,Pro917 | |
| | | Tyr29 | Pro917,Glu918,Val921 | |
| | | Leu37 | Phe927 | |
| | | Phe218 | Met598,Asn601 | |
| | | Asp224 | Leu597 | |
| | | Ala225 | Leu597 | |
| | | Pro227 | Leu597 | |
| | | Lys235 | Asn60 | |
| | | Gly236 | Gln594 | |
| | | Thr237 | Asn590 | |
| | | Asn238 | Asn590,Asn593,Gln594,Leu597 | |
| | | Ala239 | Asn590 | |
| | | Glu241 | Lys65,His589 | |

**Table S18.** Interactions between protein VIII (chain P) and hexons in the central plate region. Nomenclature and colour codes as in the previous tables.

**VIII-H4**

| | *Domain* | | | |
|---|---|---|---|---|
| P | Body | Asn16 | Glu638,Asn882 | J |
| | | Val18 | Glu638,Phe927 | |
| | | Ala23 | Asn882 | |
| | | Ile84 | Glu918 | |
| | | Asp88 | Glu918 | |
| | | Val89 | Leu597 | |
| | Neck | Gly93 | Asn593 | |
| | | Pro94 | Asn590,Asn593 | |
| | Head | Leu112 | Lys65,Ala66 | |
| | | Ser113 | Ala66 | |
| | | Gly114 | Lys65 | |
| | | Gly115 | Lys65 | |
| | | Glu175 | Asn590 | |
| | | Met176 | Asn590 | |
| | | Thr177 | Asn590,Gln594 | |
| | Neck | Lys182 | Leu597 | |
| | | Leu185 | Gln594,Met598 | |
| | | Arg186 | Leu597,Arg600, Asn601,Pro917 | |
| | | Gln188 | Asn601,Thr603 | |
| | | Gly189 | Thr603 | |
| | | Pro190 | Thr603 | |
| | Body | Ala228 | Pro878,Asn882 | |
| | | Phe240 | Asn877,Pro878,Met879 | |
| | | Glu241 | Met879 | |
| | Body | Tyr15 | Thr12 | K |
| | | Pro17 | Thr12,Pro13 | |
| | | Val18 | Arg14,Leu15 | |
| | | Thr19 | Leu15 | |
| | | Gln73 | Asp7 | |
| | | Pro75 | Thr5 | |
| | | Tyr76 | Ala3,Leu4 | |
| | | Ala77 | Ala3,Thr5 | |
| | | Ile85 | Ala3,Thr5 | |
| | | Tyr183 | Ala2 | |
| | | Leu185 | Asp32 | |
| | | Val187 | Ala2,Ala3 | |
| | | Gln188 | Glu31,Asp32 | |
| | | Gly189 | Leu4,Ser30 | |
| | | Pro190 | Leu4,Tyr28 | |
| | | Ser191 | Glu31 | |
| | | Gln192 | Pro23,Glu27 | |
| | | Glu196 | Gln16 | |
| | Neck | Val199 | Gln16 | |
| | Body | Ser201 | Thr12 | |
| | | Gln202 | Thr12 | |
| | | Phe230 | Arg59 | |
| | | Phe240 | Thr58,Arg59,Glu64 | |
| | Head | Gly106 | Met879,Asn883,His885 | L |
| | | Val107 | Asn883,His885 | |
| | | Arg109 | Met879 | |
| | | Gln111 | Gly698,Asn699 | |

**VIII-H3**

| | *Domain* | | | |
|---|---|---|---|---|
| P | Neck | Arg91 | Arg312 | I |
| | | Gly93 | Arg312 | |
| | | Pro95 | Glu918,Val921 | |
| | | Ser97 | Pro917,Glu918 | |
| | | Ala98 | Asn915,Pro917 | |
| | | Val99 | Ala602,Asn915 | |
| | | Pro101 | Ala925 | |
| | Head | Gln108 | Thr636,Glu638, Phe927,Thr929,Ala932 | |
| | | Arg109 | Ala932,Gly934,Ala936 | |
| | | Val110 | Tyr926,Phe927,Gly934 | |
| | | Gln111 | Val319,Tyr926 | |
| | | Leu112 | Ala925 | |
| | | Ser113 | Val317,Val319,Met924,Tyr926 | |
| | | Gly114 | Val317,Ala923,Met924 | |
| | | Gly115 | Ser313,Met315,Leu922,Ala923 | |
| | Head | Pro101 | Tyr17 | G |
| | | Phe102 | Leu8 | |
| | | Gln108 | Arg14 | |
| | | Val110 | Phe18 | |

(table continues in next page)

**Table S18 (continued)**

| VIII-H3 (AU1) | | | | |
|---|---|---|---|---|
| | *Domain* | | | |
| P | Body | Ala6 | Val937 | G |
| | | Pro8 | Val937 | |
| | | Thr9 | Val937 | |
| | | Val12 | Asn882,Ser884 | |
| | | Trp13 | Asn883,Ser884 | |
| | | Lys14 | Ala634,Ser884 | |
| | | Pro17 | Asp691,Ile694 | |
| | | Val18 | Ser693 | |
| | | Gln26 | Ala634,Ser884 | |
| | | Asn28 | Thr636 | |
| | | Tyr29 | Ala634,Arg635,Ala932 | |
| | | Gly30 | Arg635,Thr636, Thr929,Pro930,Ala932 | |
| | | Ala31 | Thr636,Glu638,Ala932 | |
| | | Thr32 | Phe927,Ala932, Gly934,Asn935 | |
| | | Ile33 | Ala925,Tyr926,Asn935 | |
| | | Asp34 | Ala925,Tyr926,Gly934 | |
| | | Trp35 | Asn935,Val937 | |
| | | Val36 | Asn935 | |
| | | Leu37 | Met315 | |
| | | Pro38 | Asn316,Val317,Met924 | |
| | | Gly39 | Ala923,Met924 | |
| | | Ser42 | Ala923 | |
| | | Phe43 | Arg913,Asn915, Ala923-Ala925 | |
| | | Thr60 | Asn882 | |
| | | Phe67 | Asn877,Pro878 | |
| | | Gln202 | Met879 | |
| | | Phe205 | Asn883 | |
| | | Met206 | Pro878,Met879,Asn883 | |
| | | Pro211 | Asn882 | |
| | | Ala31 | Arg14 | H |
| | | Ile33 | Arg14,Phe18 | |
| | | Phe43 | Tyr17 | |
| | | Ala46 | Tyr17 | |
| | | Ile50 | Arg14,Tyr17 | |
| | | Arg51 | Arg14 | |
| | | Arg53 | Pro13 | |
| | | Phe67 | Thr58,Arg59 | |
| | | Glu70 | Arg59 | |
| | | Ser71 | Arg59,Glu64 | |
| | | Asp72 | Thr63,Glu64,Lys65, Asp588,Asn590 | |
| | | His80 | Asp95,His589 | |
| | | Glu81 | Lys65 | |
| | | Ile84 | Asp95 | |

| VIII (AU3)-H2 | | | | |
|---|---|---|---|---|
| | *Domain* | | | |
| P | Body | Asn2 | Gln16 | D |
| | | Leu3 | Pro13,Gln16,Tyr17 | |
| | | Leu4 | Pro23,Glu27,Tyr28 | |
| | | Trp35 | Pro13 | |
| | | Val36 | Arg14,Tyr17 | |
| | | Leu37 | Phe18 | |
| | | Gly40 | Arg14 | |
| | | Ala44 | Pro13 | |
| | | Phe218 | Ser30,Asp32,Glu31 | |
| | | Asp219 | Asp32 | |
| | | Tyr11 | Asn601 | F |
| | | Lys14 | Glu918 | |
| | | Gln26 | Glu918 | |
| | | Gln27 | Asn601 | |
| | | Tyr29 | Pro917,Val921 | |
| | | Leu37 | Tyr926,Phe927 | |
| | | Pro217 | Asn601 | |
| | | Phe218 | Leu597,Met598,Asn601 | |
| | | Asp224 | Leu597 | |
| | | Ala225 | Leu597 | |
| | | Pro227 | Leu597 | |
| | | Gly236 | Gln594 | |
| | | Thr237 | Asn590 | |
| | | Asn238 | Asn590,Asn593,Gln594 | |
| | | Ala239 | Asn590 | |
| | | Glu241 | Lys65,His589,Asn590 | |

Nomenclature and colour codes as in the previous tables.

### Supplementary Figures and Legends

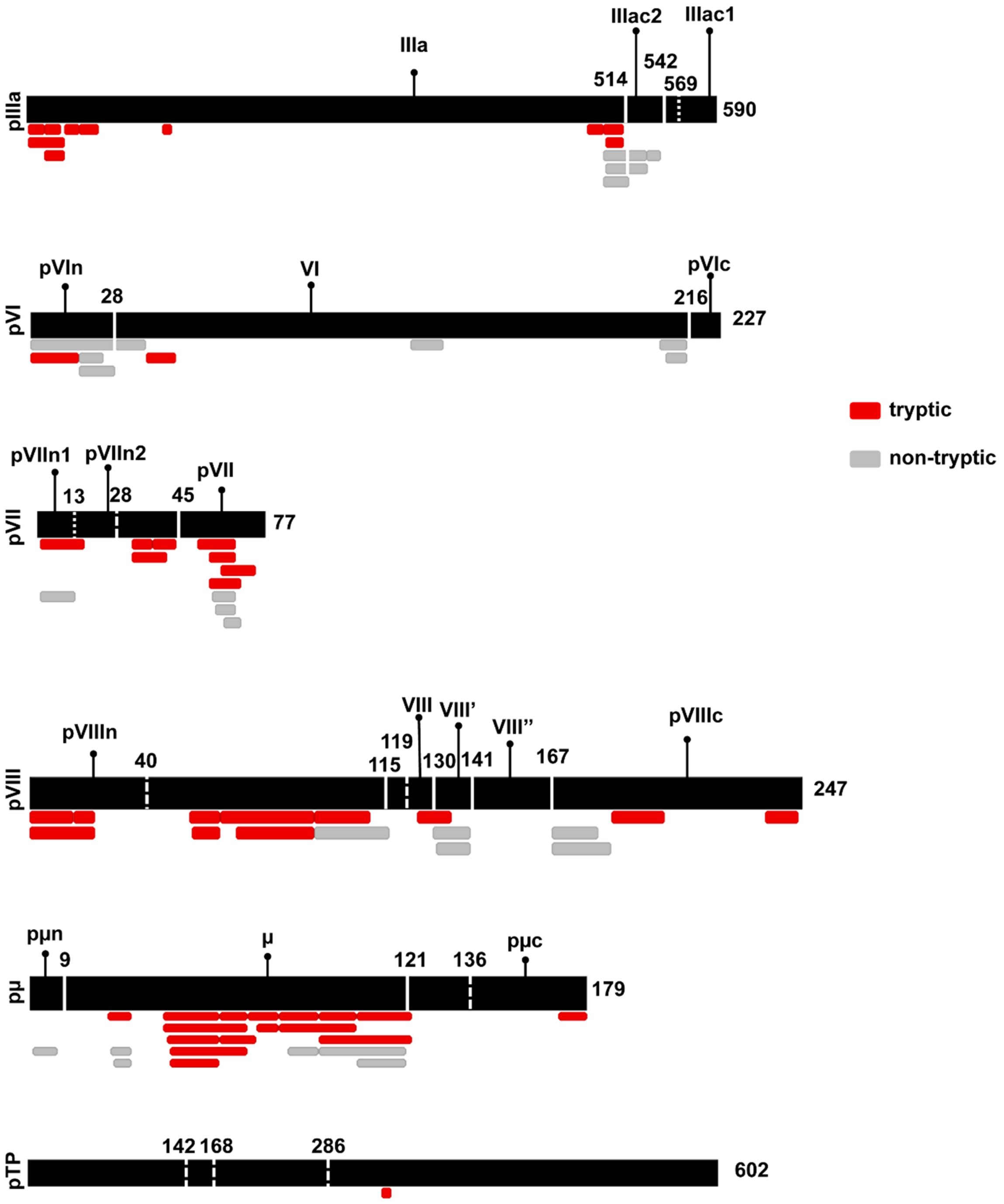

**Figure S1. FAdV-C4 proteins subject to AVP cleavage and detected in the LC-MS/MS assay.** Total lengths of immature polypeptides (in amino acids) are indicated at the right hand side. The MASCOT search was adjusted for tryptic or non-tryptic small peptides. Red bars represent fragments excised by trypsin (cuts at C-termini of Lys and

Arg, except if followed by Pro). Grey bars represent non-tryptic detected fragments, presumed to result from the endogenous AVP action. Continuous white lines indicate predicted canonical AVP sites experimentally observed in the LC-MS/MS assays. Dashed white lines indicate predicted canonical AVP sites, but not experimentally observed. Canonical AVP target sites are [MIL]xGx|G and [MIL]xGG|x (Mangel and San Martín, 2014). Dotted lines indicate non-canonical AVP sites reported in the literature for other AdVs Prefix "p" indicates precursor; "n", N-terminal excised peptide; "c", C-terminal excised peptide. L1-52/55k, which in HAdV-C5 is cleaved at multiple non-canonical sites (Pérez-Berná *et al.*, 2014), is not included here due to its complexity. The detection of peptides ending at predicted AVP cleavage sites helps confirm the universality of the AVP mechanism across the members of the *Adenoviridae* family and demonstrates that excised peptides are retained inside the capsid and not removed during assembly. Note that our structure indicates that the predicted cleavage at residue 40 in protein VIII does not occur, since we were able to continuously trace the polypeptide without interruption in that zone. Also, the central zone in pVIII is predicted to be cleaved at five sites, of which four can be confirmed in our LC-MS/MS assay but are not traced in our structure. The central region of pVIII in HAdV-C5 is cleaved at three sites (Mangel and San Martín, 2014). Differences in the cleaved sites could be significant for capsid stability, as proposed for HAdV-F41 (Pérez-Illana *et al.*, 2021).

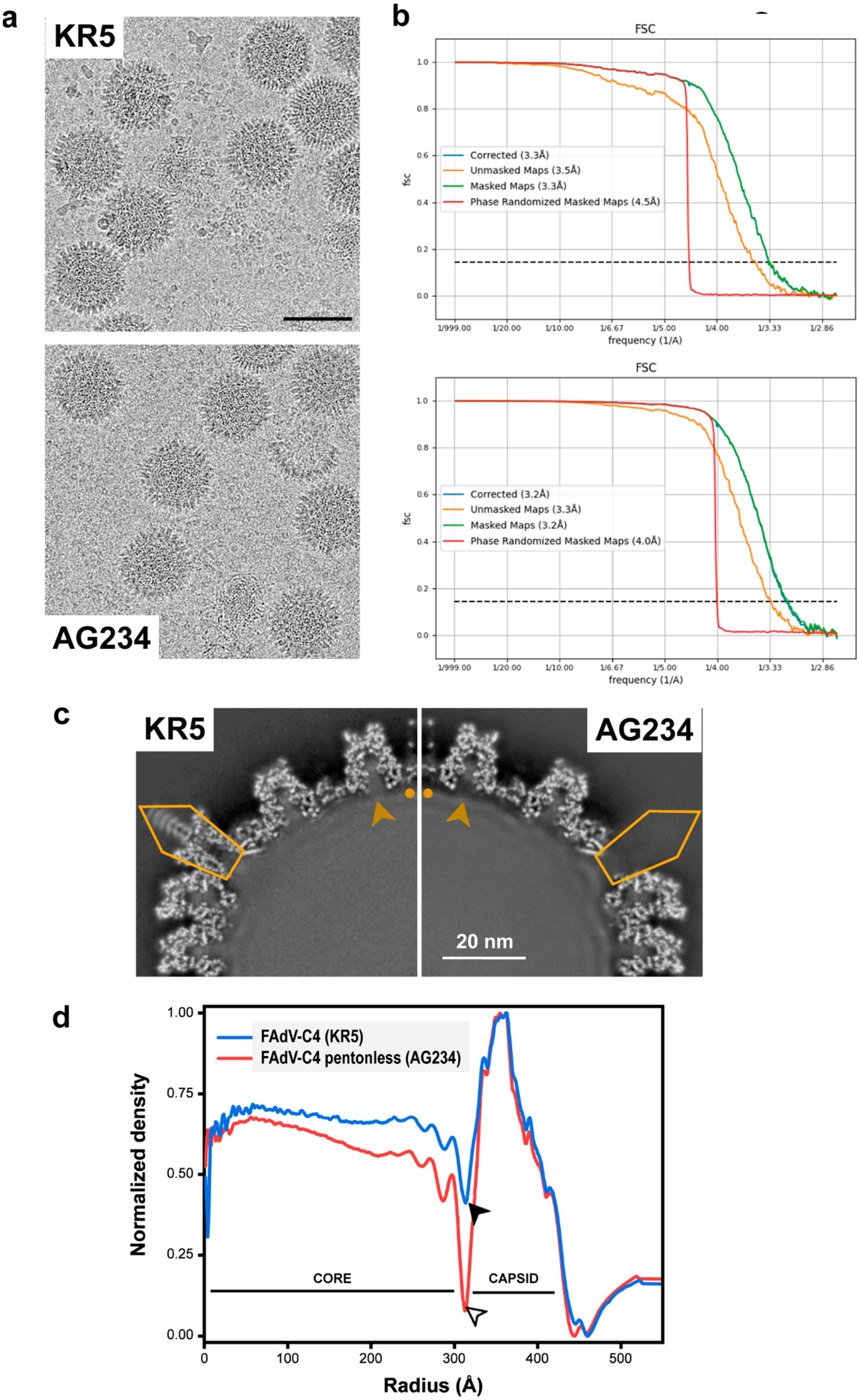
a
KR5
AG234
b
FSC
Corrected (3.3Å)
Unmasked Maps (3.5Å)
Masked Maps (3.3Å)
Phase Randomized Masked Maps (4.5Å)
frequency (1/A)
FSC
Corrected (3.2Å)
Unmasked Maps (3.3Å)
Masked Maps (3.2Å)
Phase Randomized Masked Maps (4.0Å)
frequency (1/A)
c
KR5
AG234
20 nm
d
FAdV-C4 (KR5)
FAdV-C4 pentonless (AG234)
Normalized density
CORE
CAPSID
Radius (Å)

**Figure S2. FAdV-C4 cryo-EM data and resolution. (a)** Representative area of motion-corrected micrographs for the two specimens used (FAdV-C4 strains KR5 and AG234, as indicated). The bar represents 100 nm. **(b)** Fourier Shell Correlation (FSC) curves, as provided by RELION postprocess. Note that the corrected FSC curves (blue) overlap with the curve for the masked maps (green). Resolution values at the intersection of the FSC curves with the FSC = 0.143 threshold (dotted black line) are indicated in the plot legends. **(c)** A quadrant of the central slice of the density maps viewed along a 2-fold icosahedral axis. Notice the lack of pentons in AG234. Pentagons: presence or absence of pentons. Arrowheads: presence or absence of capsid-core connections. Dots: presence or absence of RD4 (see Figure 5). **(d)** Radial average profiles of the FAdV-C4 (KR5) and FAdV-C4 pentonless (AG234) maps. The filled arrowhead indicates the presence of density in regions connecting capsid and core, while the hollow arrowhead points to a large dip for the pentonless map, indicating loss of capsid-core connections.

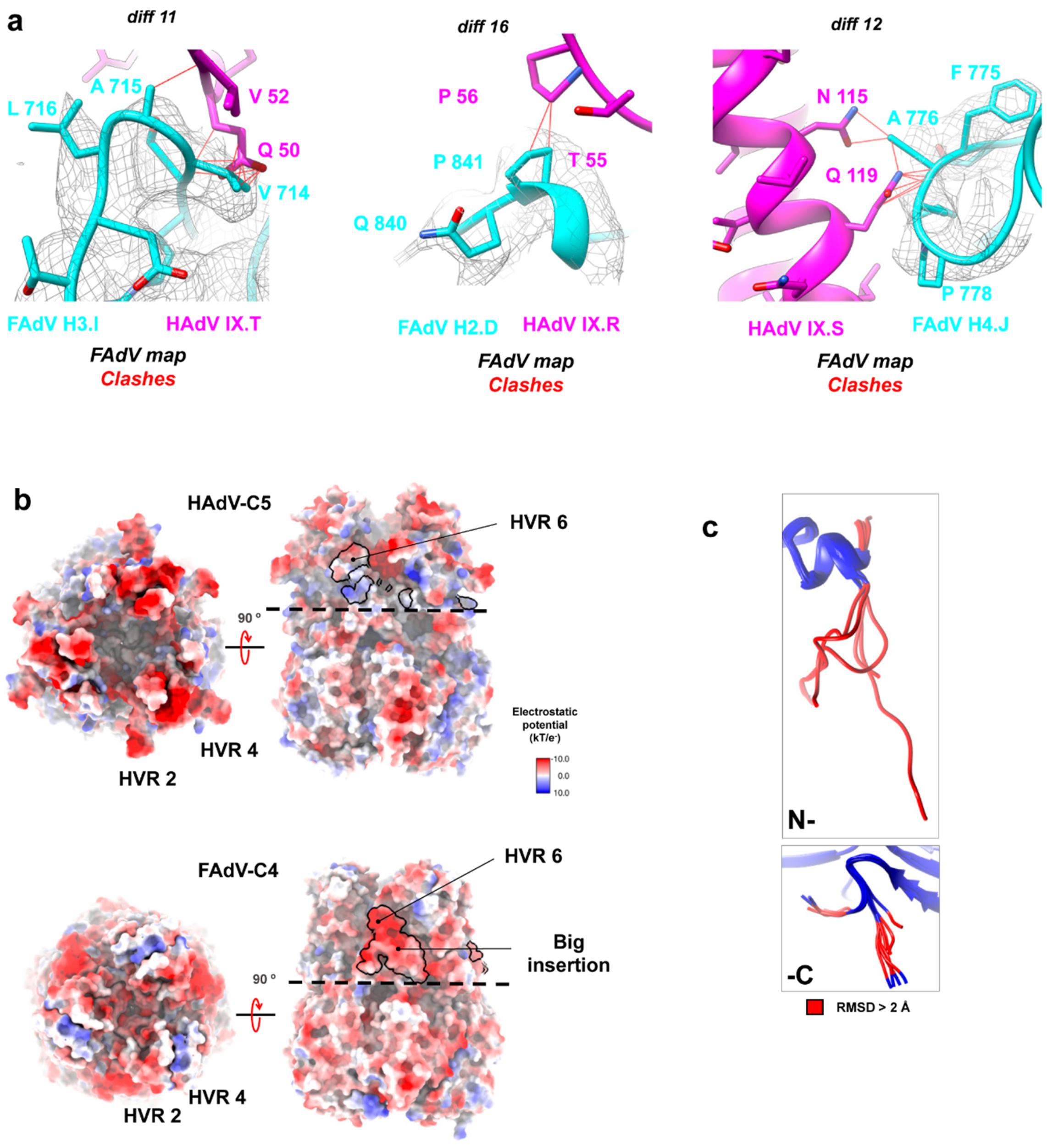

**Figure S3. Details on FAdV-C4 hexon structure and its comparison with HAdV-C5. (a)** Residues in the *pincer* region in relation to the position of protein IX in HAdV-C5. Three *diff* regions of the FAdV-C4 hexon are depicted in cyan and with the density map in grey mesh. HAdV-C5 protein IX is in pink (PDB ID: 6b1t). Hexon number (H2-4) and chain IDs are indicated. Clashes are indicated with red lines. **(b)** HAdV-C5 (PDB ID 6cgv) and FAdV-C4 hexon trimer surfaces coloured by electrostatic potential. Note that the shorter HVR2 and HVR4 in FAdV-C4 result in more compact hexon towers when compared to HAdV-C5. The big insertion (*diff 14*) and its neighbourhood (*diff 10 and 15*) are highlighted in black, and contribute to a negatively charged patch in FAdV-C4. **(c)**

Focus on the hexon termini after superposition of the twelve hexon monomers in the AU of FAdV-C4, coloured by RMSD with conserved regions in blue and residues exceeding 2 Å RMSD in red.

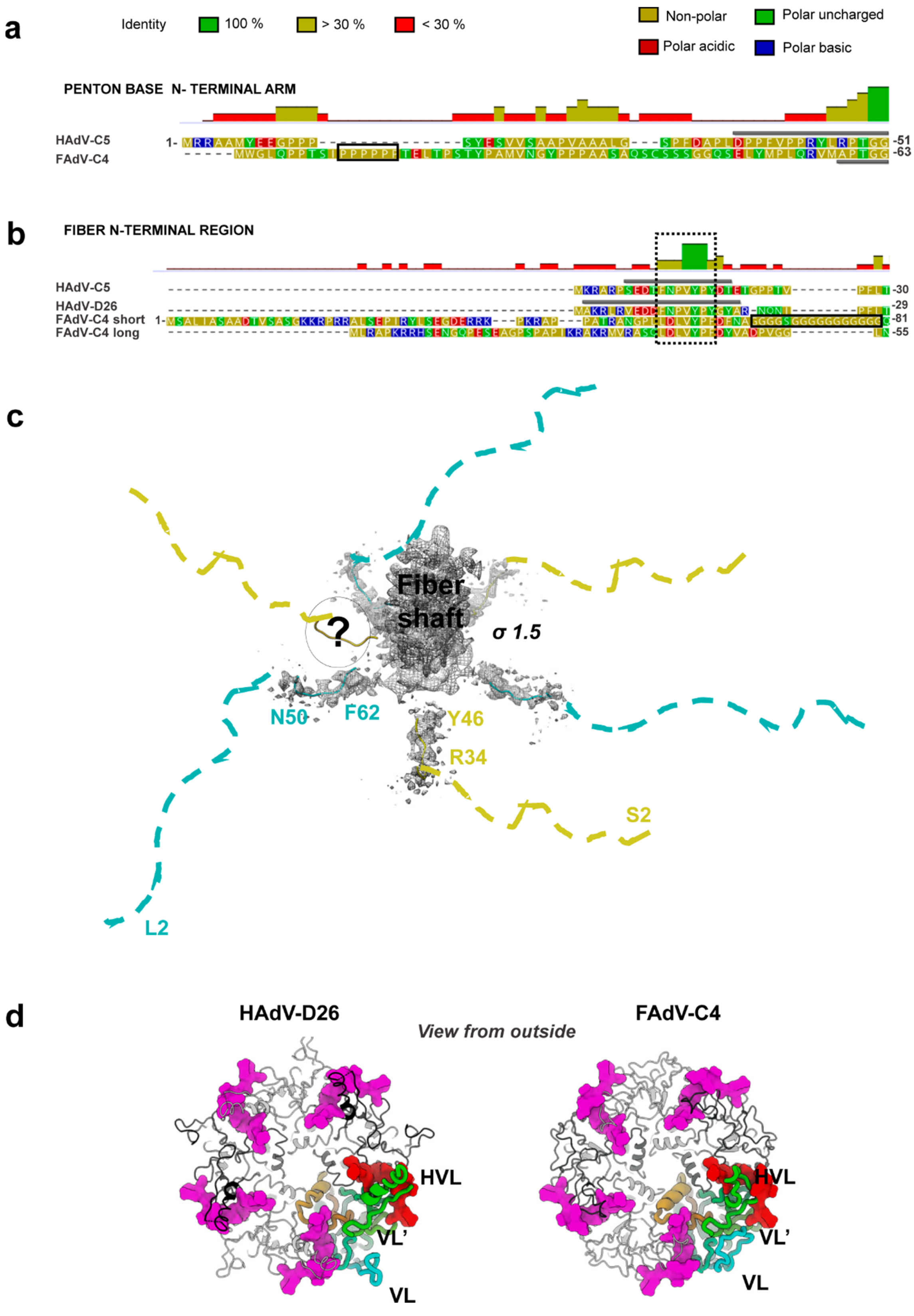

**Figure S4. Details on FAdV-C4 penton base and fibres. (a)** Sequence alignment for HAdV-C5 and FAdV-C4 penton base, focusing on the N-terminal arm. The black rectangle highlights a poly-proline stretch. **(b)** Sequence alignments for the HAdV-C5, HAdV-D26, long and short FAdV-C4 fibres, focusing on the N-terminal peptides. A black rectangle highlights the poly-glycine stretch in the FAdV-C4 short fibre. The conserved penton binding motives are indicated with a dashed rectangle. In both (a) and (b), grey bars indicate regions traced (PDB ID:6b1t for HAdV-C5 penton base, 3izo and 5tx1 for HAdV-C5 and HAdV-D26 fibres), the histograms above the sequences indicate the mean pairwise identity over all pairs in the column, and amino acids are coloured by polarity according to the colour legends at the top. **(c)** Icosahedrally averaged density for fibre(s) in FAdV-C4 (map threshold = 1.5σ), with the penton binding peptides of the long (cyan) and short (yellow) fibres fitted. The question mark indicates lack of density for one of the six N-terminal tails. There is no density to accommodate the 30 (long fibre) or 50 (short fibre) amino acids preceding the conserved penton binding motif (dashed lines). **(d)** Top view of the HAdV-D26 and FAdV-C4 penton base pentamers with the fibre N-terminal peptide modeled for HAdV-D26 overlaid. The positions of the HVL, VL and VL' are indicated for one penton base monomer (rainbow coloured). One fibre peptide is depicted in red, the other four occupying the rest of possible binding sites are in pink.

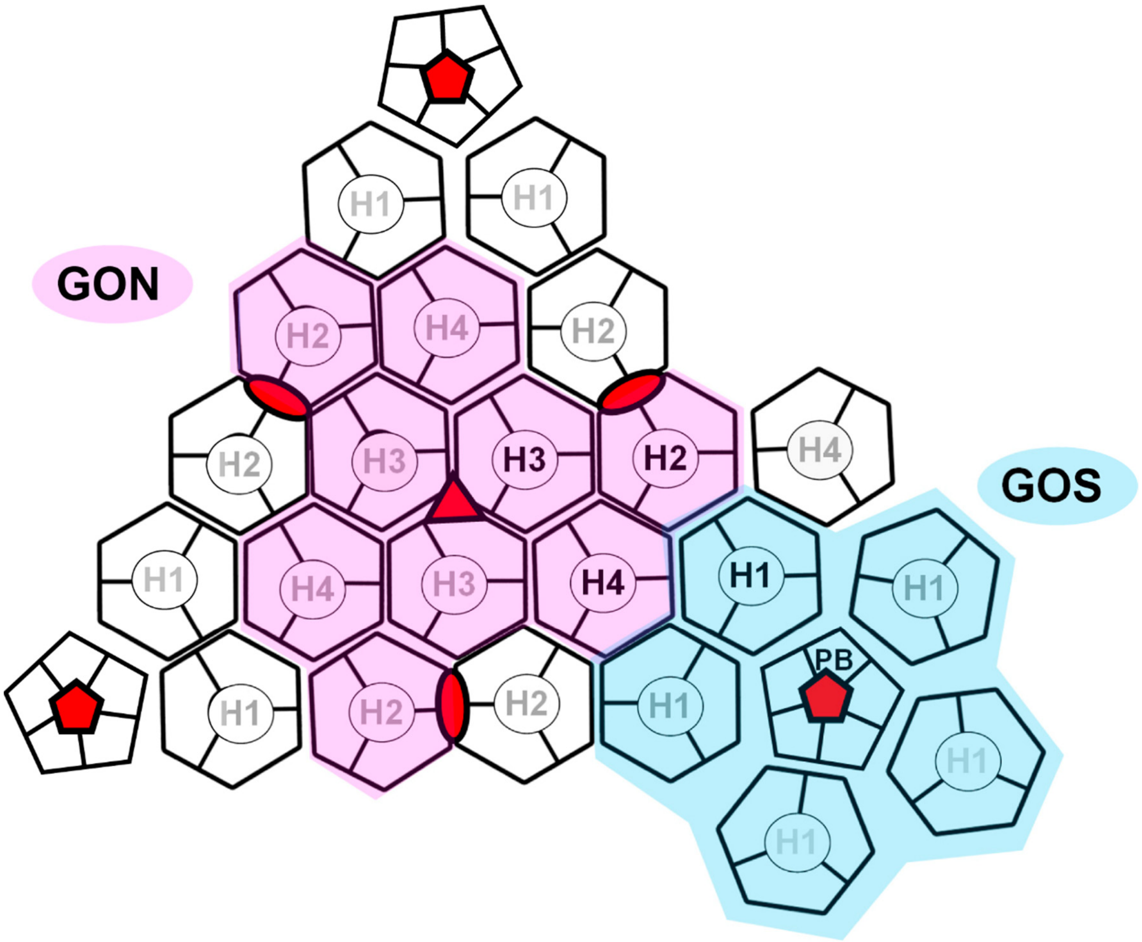

**Figure S5. Cartoon depicting geometrical features of the adenovirus capsid.** The four hexons (H1-H4) and one penton base monomer (PB) forming the icosahedral AU are labelled in black. Other hexons in the same or adjacent facets are labelled in grey. The view is from outside the capsid. Icosahedral symmetry axes are indicated with red symbols. The capsid organization can be described by two sets of tiles: (1) the Group of Six (GOS, cyan), containing one penton base pentamer and its five surrounding hexon trimers; and (2) the Group of Nine (GON, pink), containing the nine hexon trimers forming the central plate of each facet.

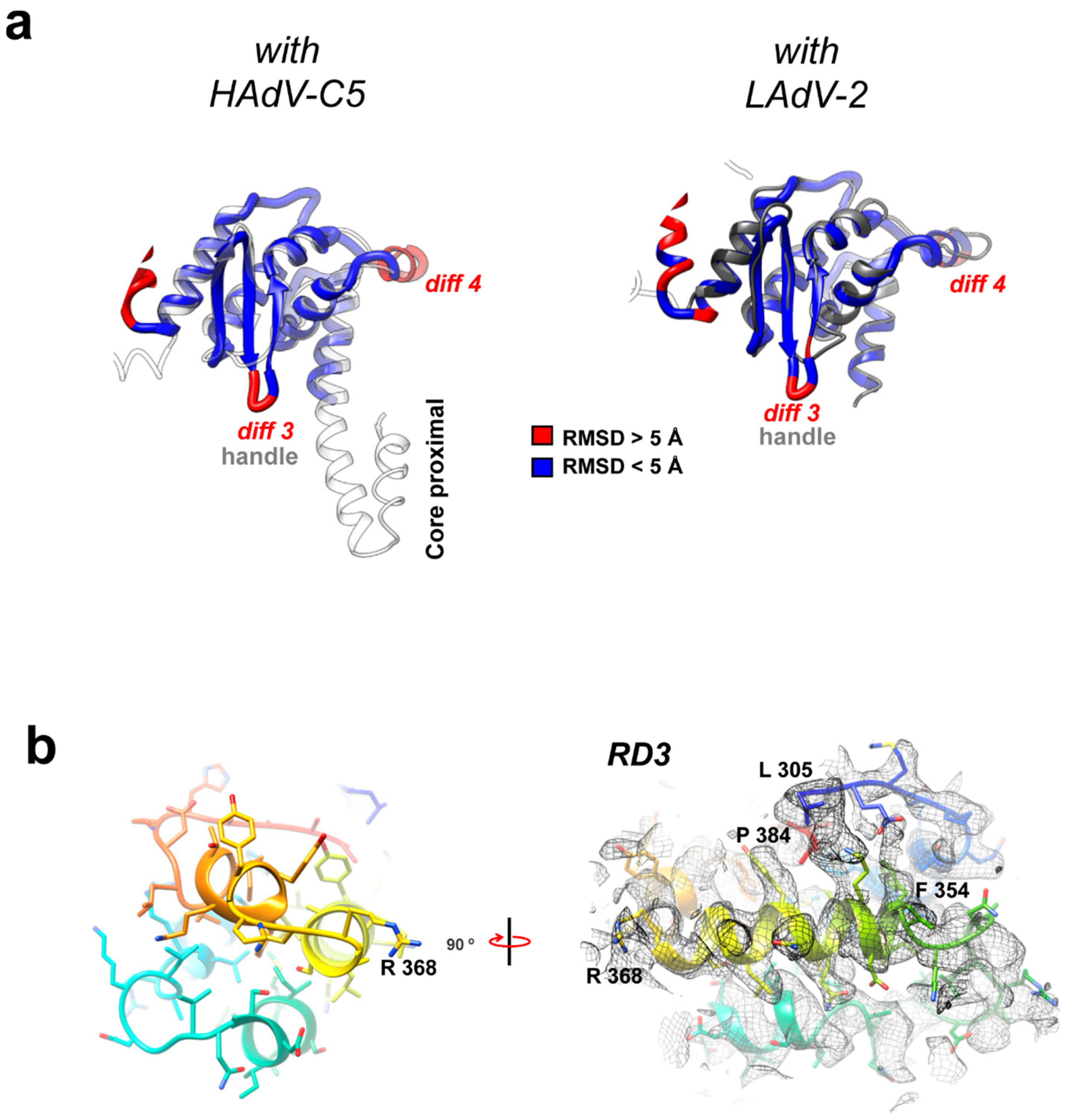

**Figure S6. Details on FAdV-C4 protein IIIa, (a)** Superposition of the VIII-binding domains of HAdV-C5 (left, white) and LAdV-2 (right, grey) with the FAdV-C4 protein, oriented as in Figure 4a, b and coloured by RMSD. **(b)** Interpretation of remnant density RD3. **Left**: Rainbow-coloured APD domain of IIIa tentatively traced as a four α-helix bundle comprising amino acids 305-384. **Right:** 90º rotated model, with the density map.

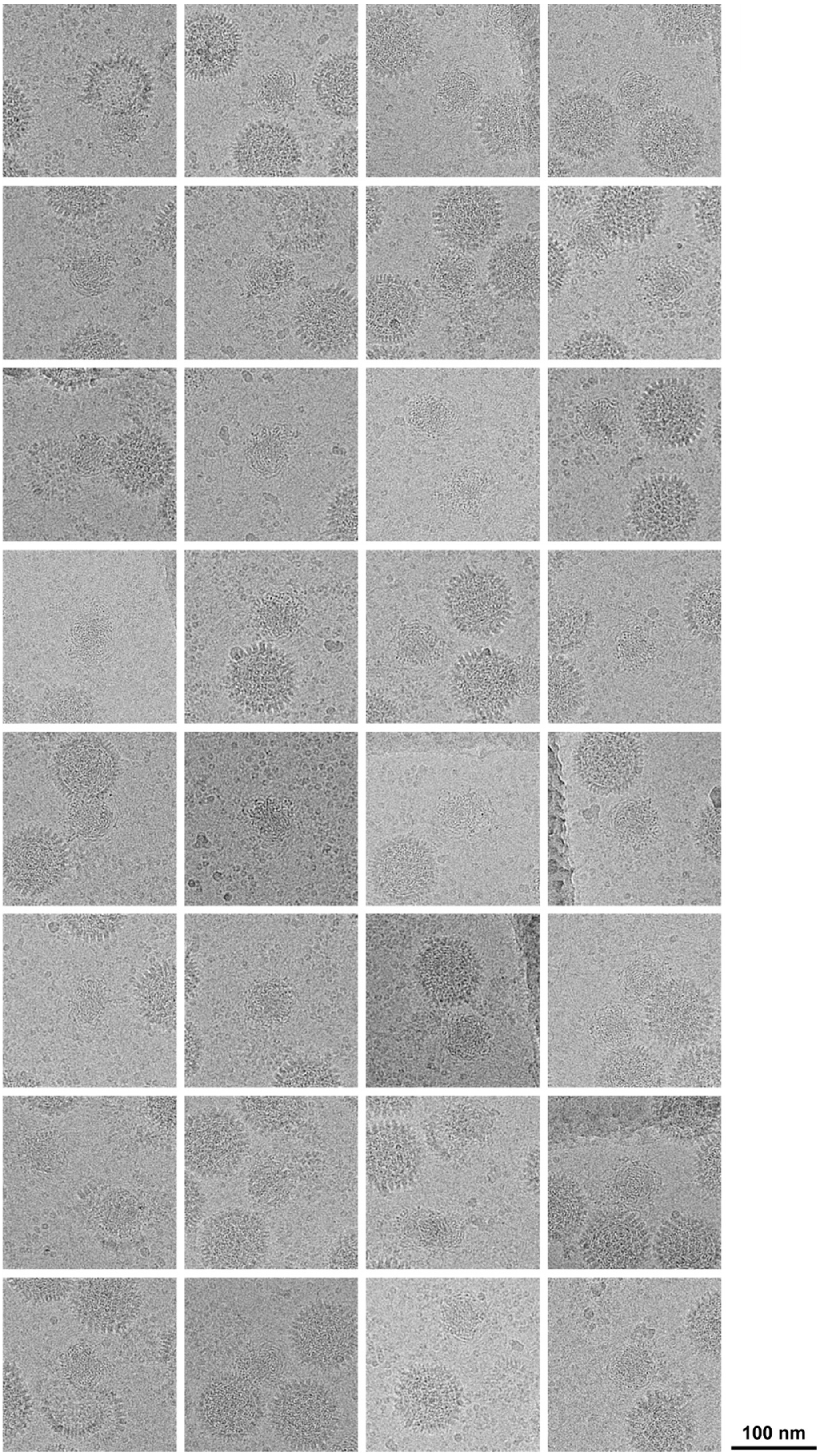

**Figure S7. Condensation of the FAdV-C4 core.** Gallery of cryo-EM images showing cores released from FAdV-C4 particles (AG243).

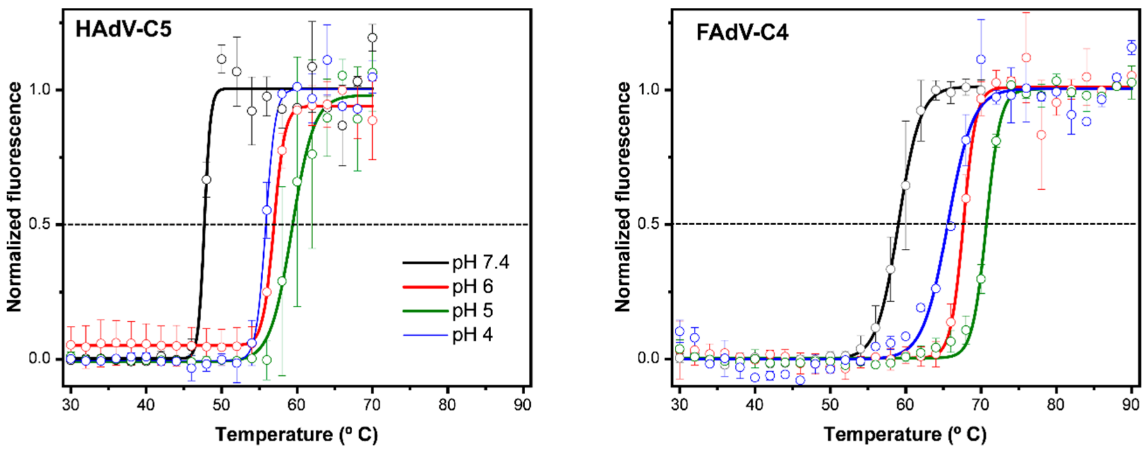

| $T_{0.5}$ ± SE (ºC) for HAdV-C5 and FAdV-C4 at different pH values. | | | | |
|---|---|---|---|---|
| | pH 7.4 | pH 6 | pH 5 | pH 4 |
| *HAdV-C5* | 47.71 ± 0.34 | 56.92 ± 0.08 | 59.34 ± 0.25 | 55.85 ± 0.11 |
| *FAdV-C4* | 59.05 ± 0.07 | 67.66 ± 0.15 | 70.71 ± 0.09 | 65.59 ± 0.35 |

*(N=2)*

**Figure S8. Combined effect of acidification and heating on HAdV-C5 and FAdV-C4 virions analysed by extrinsic fluorescence.** Open circles represent the normalized fluorescence ± STD. Continuous lines correspond to the Boltzmann sigmoid fitting to estimate the $T_{0.5}$ for each condition tested (shown in the table at the bottom). Note the increased stability of FAdV-C4 for all the conditions, compared to HAdV-C5.

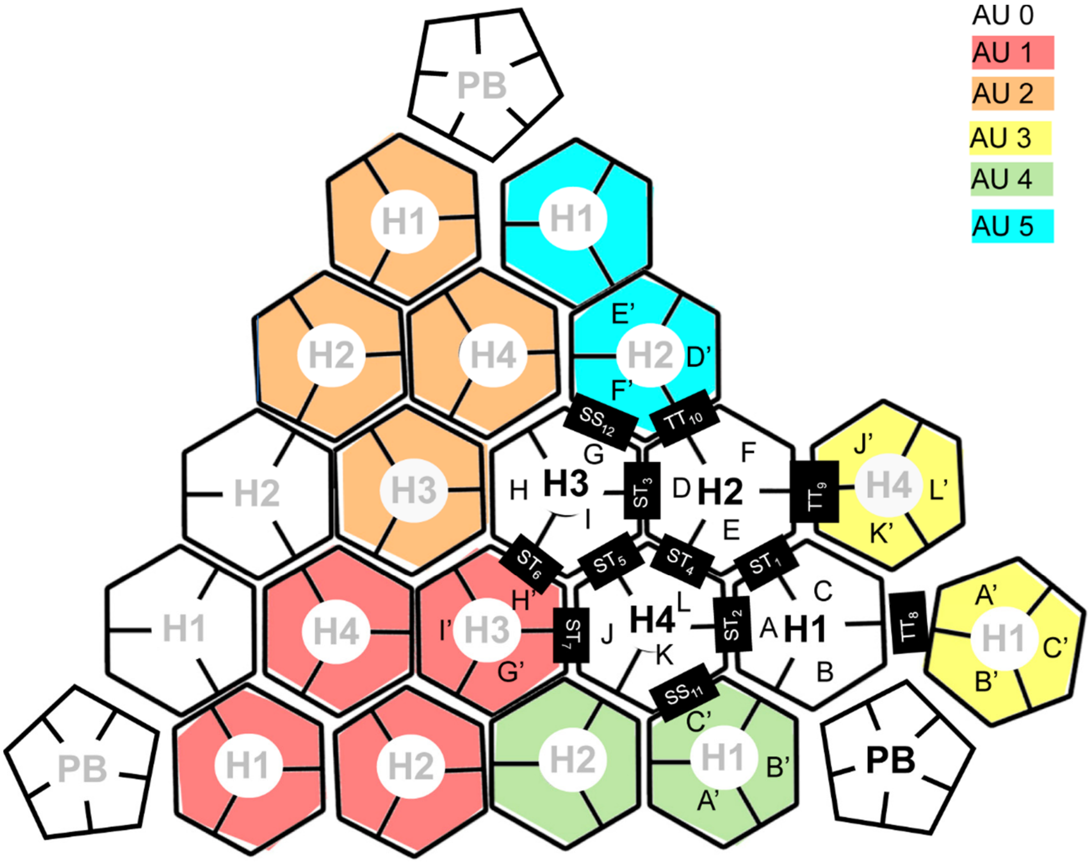

**Figure S9. Cartoon showing one AU and its neighbours.** Hexons 1-4 in one AU (AU0) are depicted in white and labelled with black text. Those in neighbouring AUs (AU1-AU5) are coloured according to the colour key at the upper right corner and labelled with grey text. Letters A-L identify different hexon chains. Interfaces between hexons are indicated in black rectangles, with S designating the facet of the hexon pseudo-hexagonal base formed by the two β-barrels in a single monomer, and T indicating the facet of the hexon pseudo-hexagonal base formed by two β-barrels belonging to two adjacent hexon monomers (Liu *et al.*, 2010). This figure accompanies the interaction data shown in Supplementary Tables S12 to S18.

**Supplementary file legend**

**File S1.** Video showing morph of proteins IIIa in HAdV-C5 (yellow, 6b1t), LAdV-2 (pink, 6qi5) and FAdV-C4 (blue, this work).
